# Cryo-EM Structure of a 95-Basepair Double-Stranded DNA Minicircle at 5.3 Å Resolution

**DOI:** 10.64898/2026.05.19.726095

**Authors:** Yukang Liu, Kyu-Yeon Lee, Yao He, Donggyun Kim, Hongjian Chang, Vadim Cherezov, Juli Feigon, Peter Z. Qin

**Author notes:** Corresponding Author: Peter Z. Qin: Department of Chemistry, University of Southern California, Los Angeles, California 90089, United States;. Department of Pharmacology and Chemical Biology, School of Medicine, University of Pittsburgh, Pittsburgh, PA 15213, United States.

## Abstract

Double-stranded DNA minicircles have been observed in a variety of biological settings and are also widely employed in biotechnology, therapeutic applications, and basic research. Here, we report a cryo-EM structure of a 95-basepair minicircle (dsMC95) at a 5.3 Å resolution. dsMC95 forms a closed ring as designed and no local deformation is observed. The two DNA strands are fully resolved, with the major and minor grooves clearly distinguishable. Analysis reveals a nine-fold periodicity in the helical twist, which corresponds to approximately 10.56 base pairs per turn. Together with groove width analysis, the data indicate that dsMC95 maintains a B-DNA configuration. The dsMC95 ring exhibits an in-plane ellipticity of 1.13 and an out-of-plane displacement of 15°, with differences in out-of-plane displacements observed between the two half-segments. The dsMC95 structure, which is the only free DNA cryo-EM structure with a resolution better than 6 Å to date, allows comparison to other structures to better understand DNA physical features such as bending. The findings advance our understanding of DNA structure under topological constraints and may inform studies of naturally occurring small circular DNA as well as the manipulation of DNA in nanotechnology applications.

## Introduction

Circular double-stranded DNA (dsDNA) forms a closed loop with no ends (1–3). Naturally-occurring circular dsDNAs are highly diverse in size and function (3–8). For example, in humans, circular double-stranded large clonal extrachromosomal DNA (ecDNA) ranges from 100 kilobases (kb) to several megabases (Mb) and represents a common mode of oncogene amplification observed across many cancer types (5, 6, 9–11). On the other hand, microDNA, which is a sub-class of extrachromosomal circular DNA (eccDNA) with lengths ranging from 80 to 400 base-pairs (bp) (3, 4), has also been found in normal tissues or as a byproduct of programmed cell death (12). Circular dsDNA has also been widely used in biotechnology, therapeutics, and basic research (1, 3). Vectors modified from naturally-occurring plasmids have been routinely used for gene manipulation and delivery. Synthetic dsDNA circles, with sizes ranging from thousands of bp to below 100 bp, have been used in nucleic acid production and therapeutics (3, 4, 13–17) as well as in DNA nanotechnology applications (18–23).

The closed-loop feature of circular DNA adds a layer of topological constraints (2, 24, 25), that in addition to the primary sequence, modulate DNA interactions with partners such as proteins (16, 26–31) and other nucleic acids (32, 33). A number of methods have been reported for synthesizing dsDNA minicircles in the range of 10s to 100s of bp (23, 26, 34–41). Conformations and dynamics of DNA minicircles have been investigated using gel electrophoresis (26, 34, 38, 42), analytical centrifugation (42), atomic force microscopy (AFM) (31, 33, 40, 43–47), electron microscopy (EM) and electron tomography (ET) (11, 12, 31, 38, 48–51), and computer simulations (42, 52–57). These studies have yielded insights into the physical properties of the DNA alone (e.g., bending and kinking (35, 38, 49, 53), curvature (57), flexibility (33, 38), and supercoiling (31, 33, 42, 44, 50)) as well as their effects on essential DNA-dependent interactions with proteins (16, 26–28, 31, 32).

Notably, EM and ET has been used to directly visualize small DNA circles (11, 12, 38, 48–51). However, published EM and ET studies of DNA circles, including those using cryo-EM and cryo-ET, are currently limited to nanometer-scale resolution and have not been able to resolve the helical structure of the dsDNA within the circles (11, 12, 38, 48–51). Indeed, although advances in cryo-EM over the past decade have enabled the determination of biomolecular structures at near-atomic resolution, cryo-EM studies on structures of DNA and RNA alone lag behind those of protein and protein-nucleic acid complexes. In particular, while at present more than 3,300 entries containing DNA have been reported in the Electron Microscopy Data Bank (EMDB), less than 20 correspond to DNA-only structures, with the best resolutions reported falling between 7.3 and 8.6 Å (58–62). In this work, we report a cryo-EM structure of a 95-bp DNA minicircle (designated as dsMC95) at a 5.3 Å resolution. dsMC95 forms a closed ring as designed, with the two strands of DNA completely resolved and major and minor grooves clearly defined. Analysis reveals a nine-fold periodicity in the helical twist. The corresponding 10.56 base-pairs per turn in helical twist, together with groove width analysis, indicate that the B-DNA configuration is maintained in dsMC95. While no severe duplex deformations such as kinking or unzipping were present, some local structural variations were observed. The dsMC95 ring shows a modest in-plane ellipticity of 1.13 and an out-of-plane displacement of 15°, with differences in out-of-plane displacement observed between the two half-segments. The dsMC95 structure represents the first DNA-only cryo-EM structure with a resolution better than 6 Å and enables direct comparison with other structures to elucidate DNA physical features such as bending. These findings expand our understanding of DNA structure under constraints, and provide a foundation for future studies of naturally occurring small circular DNA as well as for the manipulation of DNA in nanotechnology applications.

## Materials and Methods

### Synthesis of double-stranded DNA minicircle

All DNA oligonucleotides used in this work were produced via solid-phase chemical synthesis and obtained commercially (Integrated DNA Technologies, Inc., San Diego, CA). Their full sequences and other pertinent information are provided in Supporting Information (SI) sect. S1.1. Unless otherwise specified, oligonucleotides received from the vendor, which underwent desalting and passed quality control, were used without further purification. Concentration of a respective DNA was determined based on its absorbance at 260 nm (A_260_), with extinction coefficient computed from the corresponding sequence using Molbiotools (https://molbiotools.com/).

Production of dsMC95 started with enzymatic synthesis of a single-strand DNA minicircle (ssMC) from two oligonucleotides that can fold into a hairpin with a single-strand overhang (Figure 1A). Specifically, using oligonucleotides designated as A1 and A2, an A-ssMC was first synthesized (SI sect. S1.1). In a typical reaction (approximately 1 mL), A1 and A2, each at approximately 5 μM, were mixed in Annealing Buffer (20 mM Tris-HCL pH 7.5, 100 mM KCl).

**Figure 1:**
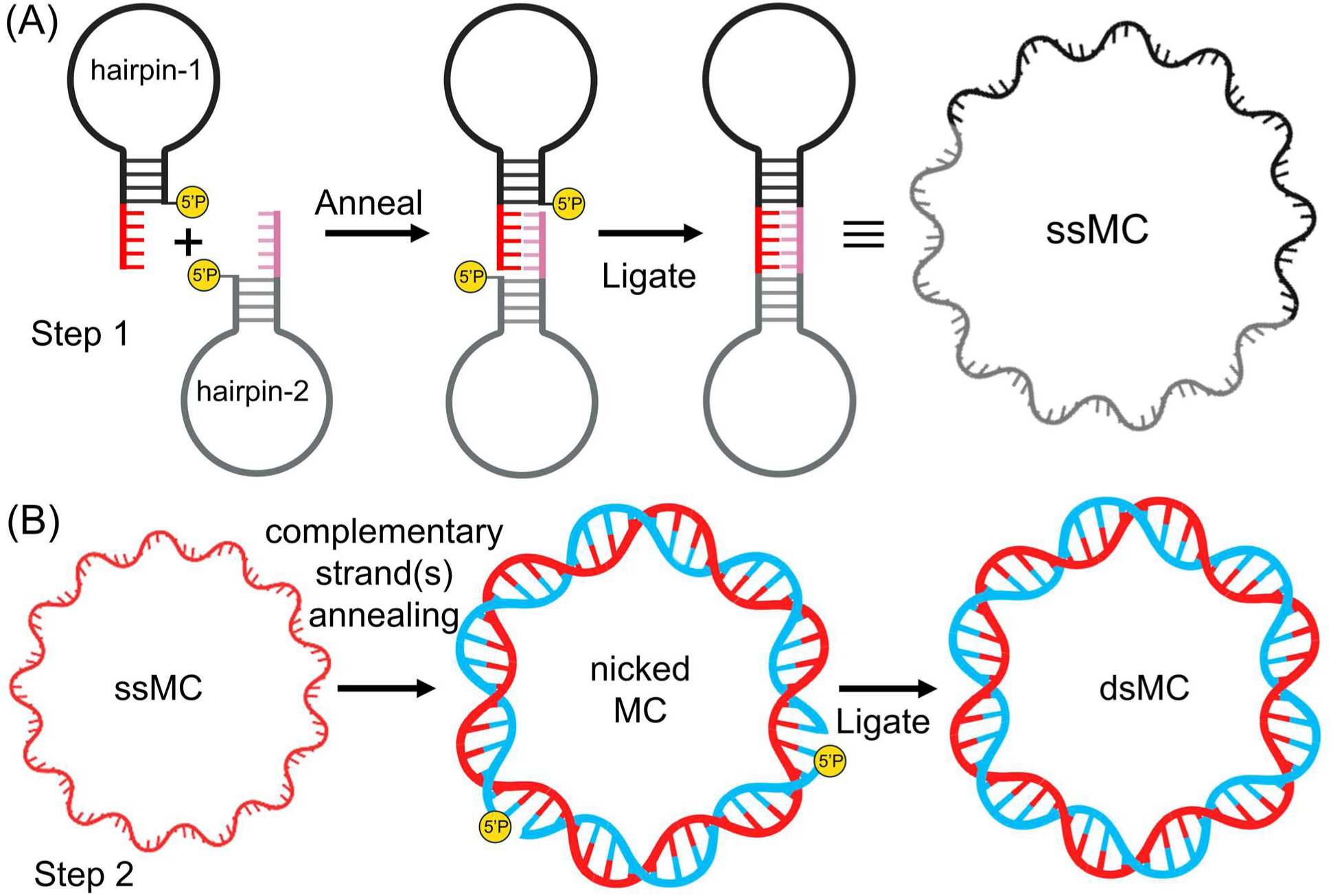
A dumbbell-mediated ligation scheme to synthesize a DNA double-stranded minicircle. (A) Step 1: Synthesis of a single-stranded minicircle. Two DNA hairpins containing complementary overhangs were annealed to form a nicked DNA dumbbell. Ligating at both termini generated a close dumbbell, which is equivalent to a single-stranded minicircle (ssMC). (B) Step 2: Synthesis of double-stranded minicircle (dsMC). A purified ssMC was annealed with one or more complementary strands to form a nicked minicircle (nicked MC). Subsequent ligation at the termini generated the fully close dsMC.

The solution was heated to 95 °C for 1 min, cooled to room temperature on the bench, and subsequently annealed at room temperature for 3 hours. Control studies showed no impact if the DNA strands were individually annealed before mixed. Upon completion of annealing, the solution was adjusted to include the T4 ligase buffer (66 mM Tris-HCl pH 7.6, 6.6 mM MgCl_2_, 10 mM DTT), 0.1 mM ATP, and T4 DNA ligase (5 Weiss U/μL, Thermo Fisher Scientific). The mixture was incubated at 16 °C for approximately 16 hours and then subjected to 10% denaturing polyacrylamide gel (29:1 acrylamide:bis-acrylamide) electrophoresis (PAGE). The A-ssMC band was excised from the gel and recovered by elution into TE (20 mM Tris-HCl pH 8.0, 1 mM EDTA), followed by filtering and concentration using Pierce™ concentrators (MWCO 10 kDa).

To produce dsMC95, an appropriate amount of A-ssMC was mixed in a 1:1:1 ratio with two 5′-phosphorylated complementary DNA strands (designated as B1 and B2, Scheme 1A, SI sect. S1.1). The mixture (generally 200 μL and contained 5 μM each of A-ssMC, B1, and B2) was adjusted to include salts specified in the Annealing Buffer, and heated to 95 °C for 1 min, then cooled to room temperature on the bench. The samples were then annealed at room temperature for 3 hours. The solution was then adjusted to include the T4 ligase buffer (66 mM Tris-HCl pH 7.6, 6.6 mM MgCl_2_, 10 mM DTT), 0.1 mM ATP, and T4 DNA ligase (5 Weiss U/μL, Thermo Fisher Scientific). The mixture was incubated at 16 °C for approximately 16 hours, then subjected to purification by 10% denaturing PAGE as described above. The purified dsMC95 was stored at -20 ℃ in TE.

### 2-amino-purine fluorescence spectroscopy

Fluorescence emission measurements of 2-aminopurine (2AP) containing DNA samples (SI sect. S1.1) were performed using a SpectraMax iD5 microplate reader (Molecular Devices, San Jose, CA). DNA samples were prepared at a final concentration of 1 μM in TE. Samples were transferred to black, flat-bottom 96-well plates to minimize background scattering, with a total volume of 100 μL per well. Excitation was set at 320 nm, and emission spectra were collected from 345 to 440 nm with a 1 nm step size. Temperature was maintained at 25 °C during data acquisition. Each measurement represents the average of three independent replicates, and baseline spectra from buffer-only controls were subtracted from the raw data. Absorbance was obtained immediately after the fluorescence measurement on a LAMBDA UV/Vis/NIR Spectrophotometer (Perkin-Elmer). Following a previously reported procedure (63), the background-corrected emission spectrum (F, obtained by subtracting the corresponding buffer emission) was normalized by A_260_ of the same sample. The resulting value at the 2AP fluorescence emission maxima, F_370_/A_260_, which is proportional to the fluorescence quantum yield of 2AP (SI sect. S.1.3), was used to assess local DNA environment at the 2AP site.

### Cryo-EM data acquisition

Purified dsMC95 was concentrated (Pierce™ concentrator, MWCO 10 kDa) to ∼12 µM in a buffer containing 20 mM Tris-HCl (pH 8.0) and 1 mM EDTA. For vitrification, approximately 3 µL samples were applied to glow-discharged UltrAuFoil^®^ R1.2/1.3 300-mesh gold grids (Quantifoil). Grids were blotted for 2–4 s at room temperature and ∼100% humidity and subsequently plunge-frozen in liquid ethane using a Vitrobot Mark IV (Thermo Fisher Scientific). Grid quality and freezing conditions were pre-screened using a Talos F200C transmission electron microscope (Thermo Fisher Scientific).

High-resolution cryo-EM data were collected on a Titan Krios cryo-electron microscope (Thermo Fisher Scientific) equipped with a K3 direct electron detector and a BioQuantum GIF energy filter (Gatan) operated at 300 kV in electron counting mode. Movies were collected at a nominal magnification of 165,000x, giving a calibrated pixel size of 0.51 Å. For the dataset reported, a total of 30,028 movies were recorded by automated data acquisition with EPU (Thermo Fisher Scientific) within a defocus range of −0.5 to −3.0 μm and a total accumulated dose of 50 e⁻/Å² per movie.

### Cryo-EM data processing

To process the dataset reported in this work, 30,028 micrographs were imported into cryoSPARC (64), where motion correction was performed using Patch Motion Correction, and the contrast transfer function (CTF) parameters were estimated using CTFFIND4 (65). Following data curation based on CTF fit resolution, ice thickness, astigmatism, and defocus, a total of 17,586 micrographs were selected for further analysis. Initial 2D classification into 10 classes (maximum resolution of 8 Å and initial classification uncertainty factor of 3) was carried out with 1,000 particles manually picked with an estimated diameter of 150 Å. The resulting 2D class averages were used as templates for template-based particle picking with a box size of 512 px and Fourier-cropped to 256 px. This procedure yielded 1,412,668 extracted particles. Three rounds of 2D classification were subsequently performed, resulting in 50 well-defined classes that were used to train a Topaz deep-learning model for particle picking. Automated particle picking was then carried out using Topaz on the 17,586 micrographs, followed by manual inspection, yielding a dataset of 530,349 particles. Three additional iterative rounds of 2D classification were conducted to eliminate poorly resolved class averages, resulting in the selection of 47 well-defined classes comprising 437,907 particles. *Ab initio* 3D reconstruction with three classes were then carried out, followed by heterogeneous refinement. This yielded a dominant class containing 347,318 particles (∼80% out of 437,907 particles). The particles in this class were then re-extracted using the original box size of 512 px to recover full resolution. Subsequent non-uniform refinement generated a final 3D density map with a gold-standard Fourier shell correlation (GSFSC) estimated resolution of 5.3 Å using the FSC=0.143 criterion.

### Atomic model generation

An atomic model of dsMC95 was manually built into the unsharpened cryo-EM density map. The model was then manually adjusted in COOT (66) and refined through multiple rounds of real-space refinement using Phenix (67). All structural figures were generated using ChimeraX (68) and PyMOL (Schrödinger, LLC). The refined dsMC95 consists of two chains, designated as i-strand and j-strand. As the circular dsMC95 has no termini, at each strand, nucleotide 95 is covalently connected with nucleotide 1. Furthermore, because the exact nucleotide identities cannot be determined using the 5.3 Å cryo-EM density map, following established precedents (29), nucleotides were designated as “dA” and “dT”, respectively, in the i- and j-strand, and the first base-pair was arbitrarily assigned within the circle.

### Analysis of radius

Analysis of dsMC95 was carried out using coordinates of the phosphate atoms. Phosphate atoms in the i-strand were designated as P_i_, with i ranging from 1 to 95 and increasing in the 5’-to-3’ direction. Phosphate atoms in the j-strand were designated as p_j_, with j ranging from 1 to 95 and increasing in the 5’-to-3’ direction. A given base-pair specified by a base-pair number (bp#) of m (i.e., bp# = m) included P_i_ with i=m and p_j_ with j=[95-(m-1)]. For example, bp#1 included P_1_ and p_95_, and bp#2 included P_2_ and p_94_.

The center-of-mass (Mc) of all the P atoms was computed by averaging the corresponding X, Y, and Z coordinates of the 190 phosphate (P) atoms. The radius of each P atom was represented by the distance between the P atom and Mc. Periodicity was analyzed using a custom-built MATLAB script to fit the radius vs. bp# data for each DNA strand according to the following equation:

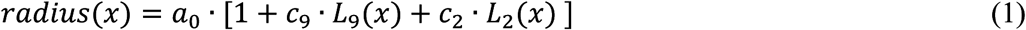

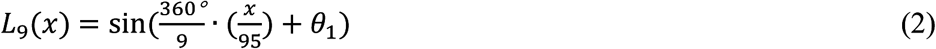

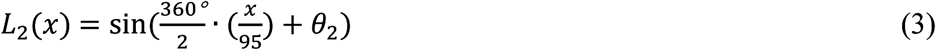

with x: the corresponding bp# of a given radius, which ranges from 1 to 95; *a*_0_: the average of the 95 individual radii; *L*_9_: a function with a periodicity of 9, with an amplitude of *c*_9_ and an initial phase of *θ*_1_; and *L*_2_: a function with a periodicity of 2, with an amplitude of *c*_2_ and an initial phase of *θ*_2_ . Fitting was carried out using a joint nonlinear least-squares regression program, nlinmultifit.m (69). The program treated *c*_9_, *c*_2_, *θ*_1_, and *θ*_2_ as variables in fitting to determine the corresponding best-fit values and their respective confidence levels. The *a*_0_ value was independently computed and not included as a variable.

### Analysis of groove widths

Following an established procedure (70), to compute the major groove width (majGW) at a given bp# of m, five distances were calculated between the i-strand P_i’_ (i’=m-2) and the j-strand p_j’_, with j’=[95-(m-1)]-n and n = 1, 2, 3, 4, 5. The smallest value among these five distances was chosen as majGW. To account for the lack of termini with the circular dsMC95, if i’=(m-2) ≤ 0, i’ was set to 95+(m-2); and if j’=[95-(m-1)]-n ≤ 0, j’ was set to {[95-(m-1)]-n}+95. Furthermore, the mid-point between the P_i’_ and p_j’_ phosphates used to calculate majGW was computed, and the distance between this mid-point and the dsMC95 center-of-mass, R_major_, was computed to assess positioning of the corresponding major groove with respect to the interior or exterior of the DNA circle.

Similarly, to compute the minor groove width (minGW) at a given bp# of m, five distances were calculated between i-strand P_i’’_ (i’’=m+1) and j-strand p_j’’_, with j’’=[95-(m-1)]+n and n = 1, 2, 3, 4, 5. The smallest value among these ten distances was chosen as minGW. If i’’=(m+1) > 95, i’’ was set to (m+1)-95; and if j’’=[95-(m-1)]+n > 95, j’’ was set to {[95-(m-1)]+n}-95. Furthermore, the mid-point between the P_i’’_ and p_j’’_ phosphates used to calculate minGW was computed, and the distance between this mid-point and the dsMC95 center-of-mass, R_minor_, was computed to assess positioning of the corresponding minor groove with respect to the interior or exterior of the DNA circle.

### Analysis using the internal coordinate of the double-stranded minicircle

Using the pdb file of dsMC95, which describes the atoms using the default laboratory coordinate, the mid-point of a given base-pair (mP) was computed by averaging the coordinates of the pair of P atoms assigned to the same base-pair. The base pair vector 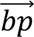, which orientates at Mc and terminates at a given mP, was computed as:

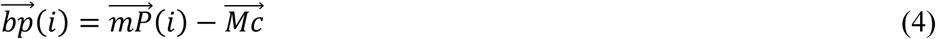

with 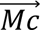 being the vector of the center-of-mass (Mc, defined above in “Analysis of radius”) expressed in the laboratory coordinate, and 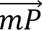(*i*) being the vector of the given mP expressed in the laboratory coordinate.

The internal coordinate of the double-stranded minicircle, designated as the dsMC coordinate, was defined with its origin set at Mc. The X-axis (*̂x*) was set at one of the 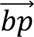 vectors designated as 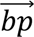(a).

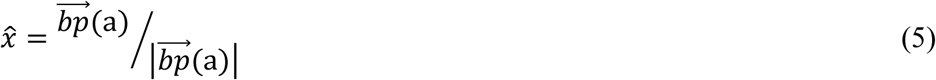

the XY-plane was defined by 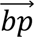(a) and another 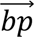 vector designated as 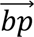(b), so that the Z-axis of the dsMC coordinate (̂*z*) was calculated as the cross product of 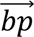(a) and 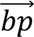(b):

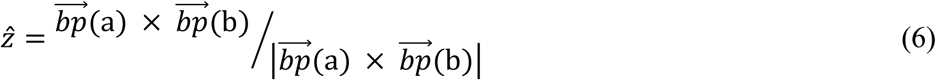

the Y-axis of the dsMC coordinate (̂*y*) was then calculated as the cross product of ̂*z* and ̂*x*:

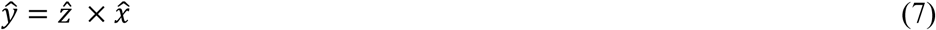

Upon establishing the three orthogonal axes, coordinates of each mP in the dsMC coordinate were computed as:

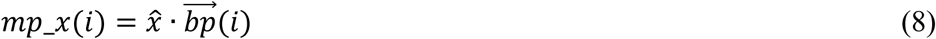

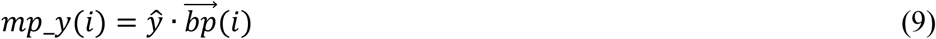

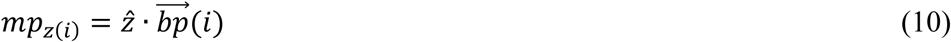

To analyze displacements of mP from the XY-plane, displacement angles were computed from the Z-coordinates of mP of the dsMC coordinate (i.e., mp_z(i), eq 10) according to:

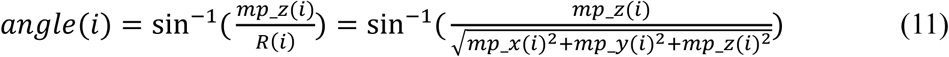

with R being the radius of a given mP. Values of *angle*(*i*) span a range of (–90°, +90°), with 0° to +90° corresponding to *mp*_*z* ≥ 0, and –90° to 0° corresponding to *mp*_*z* < 0. To analyze features at the XY-plane, projections of mP at the XY-plane of the dsMC coordinate were fit to an ellipsoid described as:

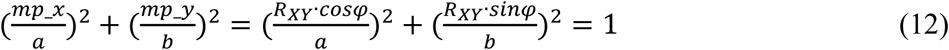

With “a” and “b” being, respectively, the long- and short-axis of the ellipsoid, mp_x and mp_y being the respective X and Y (Cartesian) coordinates of a given mP in the dsMC coordinate (eq.s 8 and 9), R_XY_ being the radius of mP at the XY-plane, and φ being the polar angle with respect to the X-axis.

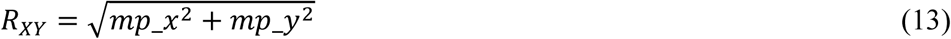

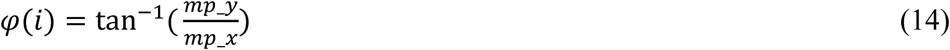

Note that values of φ span a range of (–180°, +180°), with 0° to +90° corresponding to *mp*_*x* ≥ 0 and *mp*_*y* ≥ 0; +90° to +180° corresponding to *mp*_*x* < 0 and *mp*_*y* ≥ 0; –90° to 0° corresponding to *mp*_*x* > 0 and *mp*_*y* <0; and –180° to –90° corresponding to *mp*_*x* < 0 and *mp*_*y* <0.

Based on eq. 12, the mP data set [φ(i), R_XY_(i)] were fitted to:

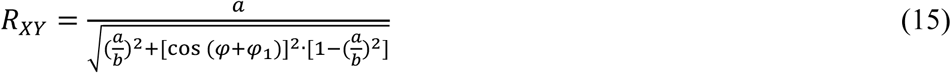

with φ_1_ representing an angle included to account for possible rotation of the ellipsoid major axis with respect to the dsMC coordinate. Fitting was carried out using a nonlinear least-squares regression MATLAB program, nlinmultifit.m (69). a, b, and φ_1_ were treated as fitting variables to determine their best-fit values and the respective confidence levels.

## Results

### Synthesis and biochemical characterization of a 95 base-pair double-stranded minicircle

A dumbbell-mediated ligation method (23) was adapted to synthesize closed and nicked DNA minicircles (Figure 1, SI sect. S1.1). A single-stranded DNA minicircle (ssMC) was first produced by ligating two hairpins containing short and complementary sticky ends (Figure 1A). The resulting dumbbell-shape ssMC was then hybridized with two linear complementary strands, and subsequent ligation of the two nicks yielded a closed double-stranded minicircle (dsMC) (Figure 1B). In addition, a single nicked minicircle can also be generated by hybridizing a full-length linear complementary strand to the ssMC while omitting ligation.

Following this scheme, a 95-bp minicircle (dsMC95) was synthesized using hairpin fragments of 48 and 47 nucleotides (SI, sect. S1.1). While results reported here focus on the free minicircle, the dsMC95 sequence was also designed for studying DNA recognition by CRISPR-Cas9 and included a Cas9 targeting sequence. Furthermore, when necessary, a fluorescence tag was incorporated for tracing the DNA products, and a 2AP was substituted at a specific site for monitoring local environment of the DNA (SI sect. S1.1, also see Results later).

dsMC95 was biochemically characterized by several approaches (Figure 2, SI, sect. S1.2). Denaturing PAGE analysis clearly revealed a single band for the expected dsMC95 species (Figure 2A, lane 3), which migrated slower than that of ssMC and the 95-nt linear strand(s) (Figure 2A). Furthermore, the nicked minicircle (nicked MC95) denatured into the respective ssMC and linear strand (Figure 2A, lane 2), while dsMC95 did not denature (Figure 2A, Lanes 2 and 3). This is consistent with previous reports (26) and provides an unambiguous test to distinguish the fully closed dsMC95 from those containing nick(s). The denaturing PAGE results were supported by additional biochemical analysis (SI, sect. S1.2). Overall, the synthesis yielded predominantly dsMC95 with minimal off-pathway species such as linear concatemers (SI, sect. S1.2.1).

**Figure 2:**
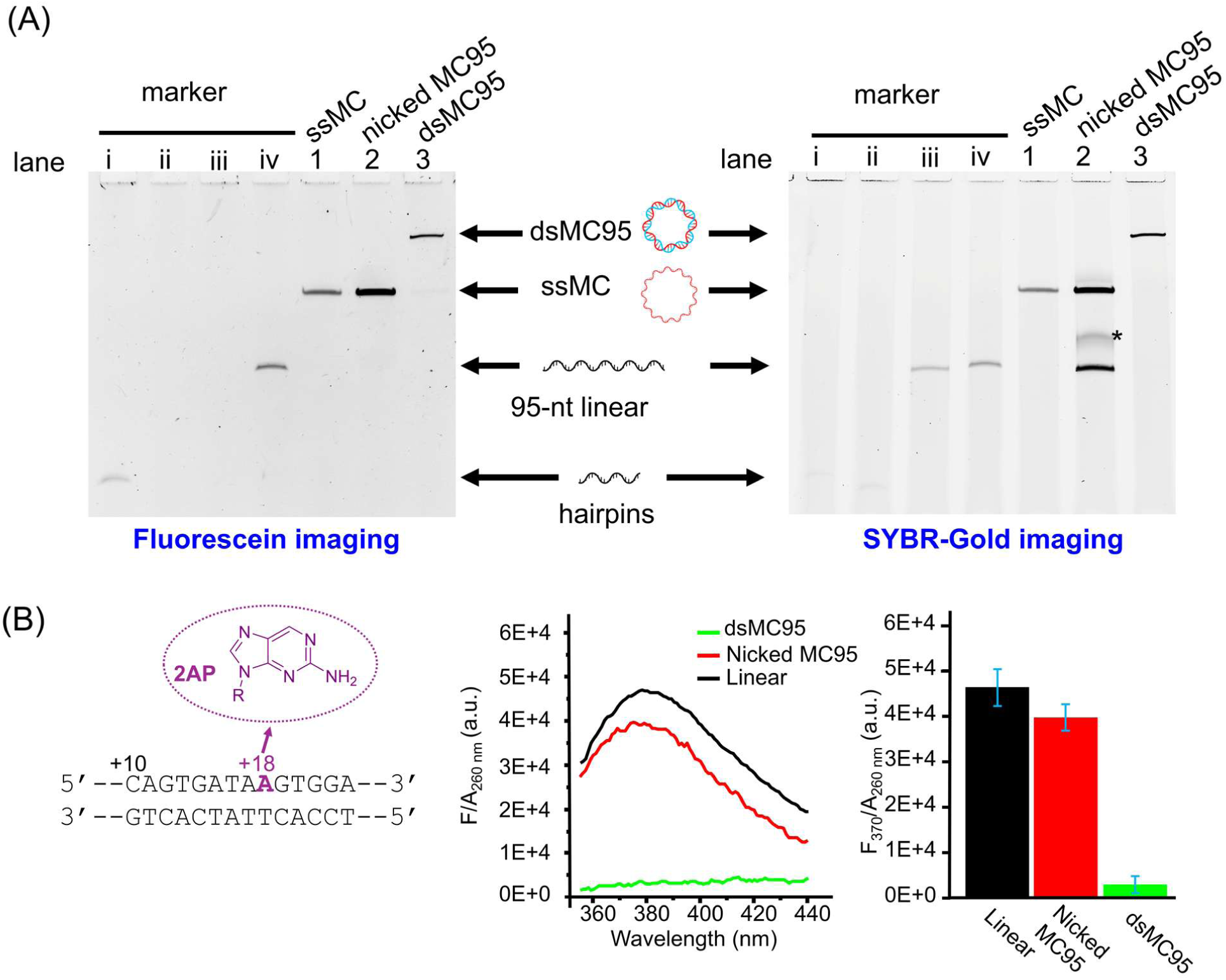
Characterization of dsMC95. (A) Biochemical characterization using denaturing PAGE. The samples were visualized by fluorescence imaging (left) and SYBR-gold dye staining (right). Lane 1 - purified ssMC (A-ssMC, see Figure S1B caption); lane 2 - nicked MC formed by annealing a ssMC (A-ssMC) with a 95-nt complementary strand (B-linear, see Table S1); lane 3 - purified dsMC95 (see Figure S1B). Marker lanes correspond to (i) a 48-nt DNA containing a fluorescent label (“A1-40-FAM”, Table S1); (ii) a 47-nt DNA without fluorescent label (“A2”, Table S1); (iii) a 95-nt linear DNA strand without fluorescent label (“B-linear” Table S1); (iv) a 95-nt linear DNA strand containing a fluorescent label (A-ssMC synthesized using “A1-40-FAM” and “A2”). “*” marks a band that may represent a minor aberrant species among the B-linear DNA. See additional information in SI sect. S1.1 and S1.2. (B) Local base-pair environment characterized by 2AP fluorescence emission. Shown on the left is the DNA construct for 2AP measurements, with the 2AP position shown in red, and its chemical structure shown in the inset. Shown in the middle is the F/A_260_ spectra measured for the same DNA sequence in different topological forms, with “nicked MC95” formed by annealing A-ssMC and B-linear strands, and “Linear” formed by annealing A-linear and B-linear strands (Table S1). Shown on the right is the comparison of F_370_/A_260_ values, with average and standard deviation (error bar) obtained from three measurements. See additional information in SI sect. S1.3.

dsMC95 was further assessed using a previously developed 2AP fluorescence assay (63). A 2AP was site-specifically substituted within the DNA duplex (Figure 2B, left), 2AP fluorescence was measured in the context of dsMC95, nicked MC95, and linear duplex (Figure 2B, middle). The F_370_/A_260_ values, which are proportional to the quantum yield of 2AP (Methods, SI sect. S.1.3) and report on the local DNA environment (for example, base stacking (63)), were obtained (Figure 2B, right). The F_370_/A_260_ value of the nicked MC95 was approximately 85% of that of the linear duplex (Figure 2B), indicating minor difference in the local environment at the 2AP incorporation site between these two constructs. In contrast, with dsMC95, no clear 2AP fluorescence peak was observed, and the F_370_/A_260_ value was approximately 8.5% of that of the linear duplex (Figure 2B). This pronounced reduction in 2AP fluorescence confirms the successful synthesis of dsMC95. It also indicates that the closed minicircle presents a very different local environment as compared to that of the nicked circle and the linear duplex.

### Single-particle cryo-EM analysis of dsMC95 reveals a mostly circular and flat DNA ring

High-resolution cryo-EM data were acquired to investigate structural features of dsMC95 (see Methods). For the dataset reported here, approximately 18,000 micrographs were selected for analysis, in which circular-shaped DNA particles were readily identifiable (SI sect. S2, Figure S5). Approximately 438,000 DNA particles were selected for 2D classification (SI sect. S2, Figure S5), and three distinct classes were identified during ab initio 3D reconstruction (SI sect. S2, Figure S5). Differences between the three classes were minimal, and one of the classes, designed as class 0, accounted for approximately 80% of the total particles (SI sect. S2, Figure S5). Class 0 was subsequently selected for further 3D refinement, resulting in an EM map with a resolution of 5.3 Å (Figure 3A; SI sect. S2, Figures S5 and S6). Validation analyses confirmed that the majority of dsMC95 orientations were well-sampled, and the gold-standard Fourier shell correlation (GSFSC) resolution estimate is reliable (SI sect. S2, Figure S6).

**Figure 3:**
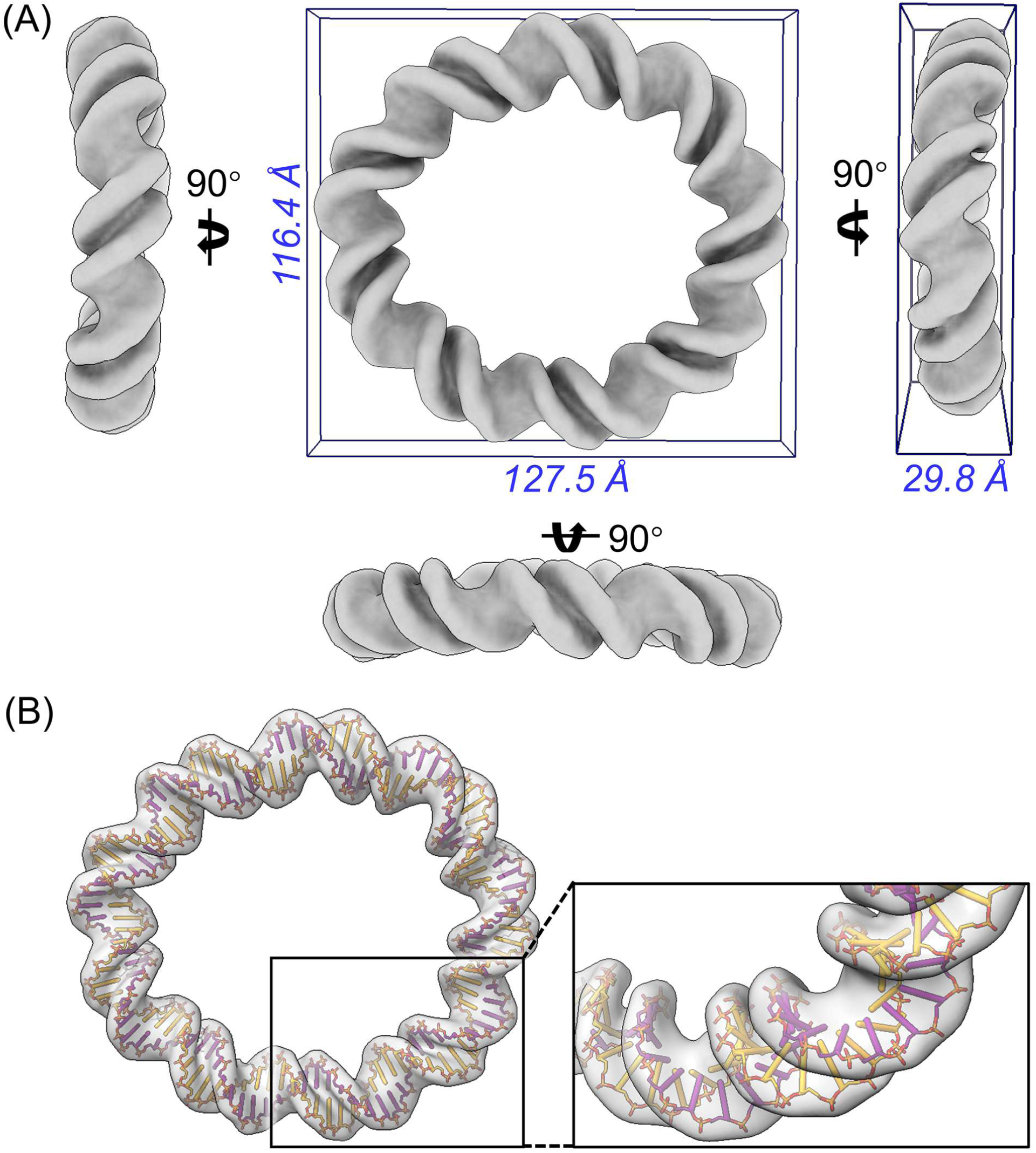
cryo-EM analysis of dsMC95. (A) Unsharpened cryo-EM map viewed from different perspectives. The map is shown using ChimeraX at a contour level of 0.018. The box shows the estimated dimensions obtained using the bounding box function in ChimeraX. (B) A full atomic model fitted within the unsharpened map (contour level 0.018.). Chain-i and chain-j of the DNA strands are colored orange and purple, respectively. Sticks represent the phosphate backbone of the DNA, bars represent the remaining portion of the nucleotides. A zoom-in view of the boxed segment is shown on the right. Additional information on cryo-EM data collection and processing is presented in SI sect. S2, with statistics summarized in Table S3. See also SI Movies S1 and S2.

The 5.3 Å map of dsMC95 shows that the DNA forms a closed ring as designed (Figure 3, SI Movie S1). The backbones of the two DNA strands can be confidently resolved, and the major and minor grooves are clearly distinguishable (Figure 3). When viewed from the top, the DNA ring is mostly circular, with dimensions estimated as approximately 128 Å × 116 Å (Figure 3A, top view). When viewed from the side, the width is estimated to be approximately 30 Å (Figure 3A, side view). This is larger than the B-DNA helix width of ∼20 Å, and indicates a certain degree of torsional stress that keeps the DNA ring from being completely flat (planar). Importantly, both strands of the DNA can be traced continuously around the entire circle, and the map shows no locally disordered segments (Figure 3). This indicates that no severe DNA duplex disruptions, such as kinking or unzipping, are evident in this 5.3 Å resolution map.

### Global Features of DNA helices in dsMC95

A full atomic model of dsMC95 was built based on the 5.3 Å map (Figure 3B, SI Movie S2). However, the map resolution was insufficient to resolve individual base-pair(s). Consequently, the DNA base pairs were designated generically as “dA:dT”, and the first base pair was assigned arbitrarily within the circle (see Methods). Further analysis of the dsMC95 conformation was performed using only the coordinates of the 190 phosphate (P) atoms of the two 95-nt DNA strands.

To analyze the conformation of dsMC95, a previously reported procedure (55) was adopted to calculate the distances between each P atom and the center-of-mass (Mc) of all P atoms (see Methods). These distances were used to represent the radius of each nucleotide from the center of the DNA ring (Figure 4A). The plots of radius vs. corresponding base pair number (bp#) show clear oscillations for the individual i- and j-strands (Figure 4B), with smaller radii corresponding to phosphate groups oriented toward the center of the ring (“inside”), whereas larger radii corresponding to phosphates directed outward (“outside”). For each strand, the oscillation can be fitted by a sum of two periodic functions (Figure 4B, “measured” vs. “simulated”; SI sect. S3.1, Figure S7). The major oscillation, L_9_ (eq. 2, Methods), shows a period of 9 (Figure 4B, 4C). This reflects the helical twist of the phosphates along the DNA strand, and the corresponding periodicity of 95/9 ∼ 10.56-bp/turn fits well with that of a B-DNA (35). Furthermore, the amplitude coefficients of L_9_ (c_9_, eq. 2, Methods) have nearly identical values but an opposite sign between the i- and j-strands, indicating an 180° phase shift (Figure 4B, 4C). This reflects the antiparallel organization of the duplex, with the two strands alternating in their respective orientations towards the ring center approximately every 5-bp. In addition, a secondary oscillation with a period of 2 (L_2_, eq. 3, Methods) is evident in the radius plots for both strands (Figure 4B), with nearly identical phases (i.e., same sign for the amplitude coefficients c2, Figure 4C). This likely reflects a deviation of dsMC95 from a perfect circle, which can already be observed from the EM map (Figure 3A) (see further analysis below).

**Figure 4:**
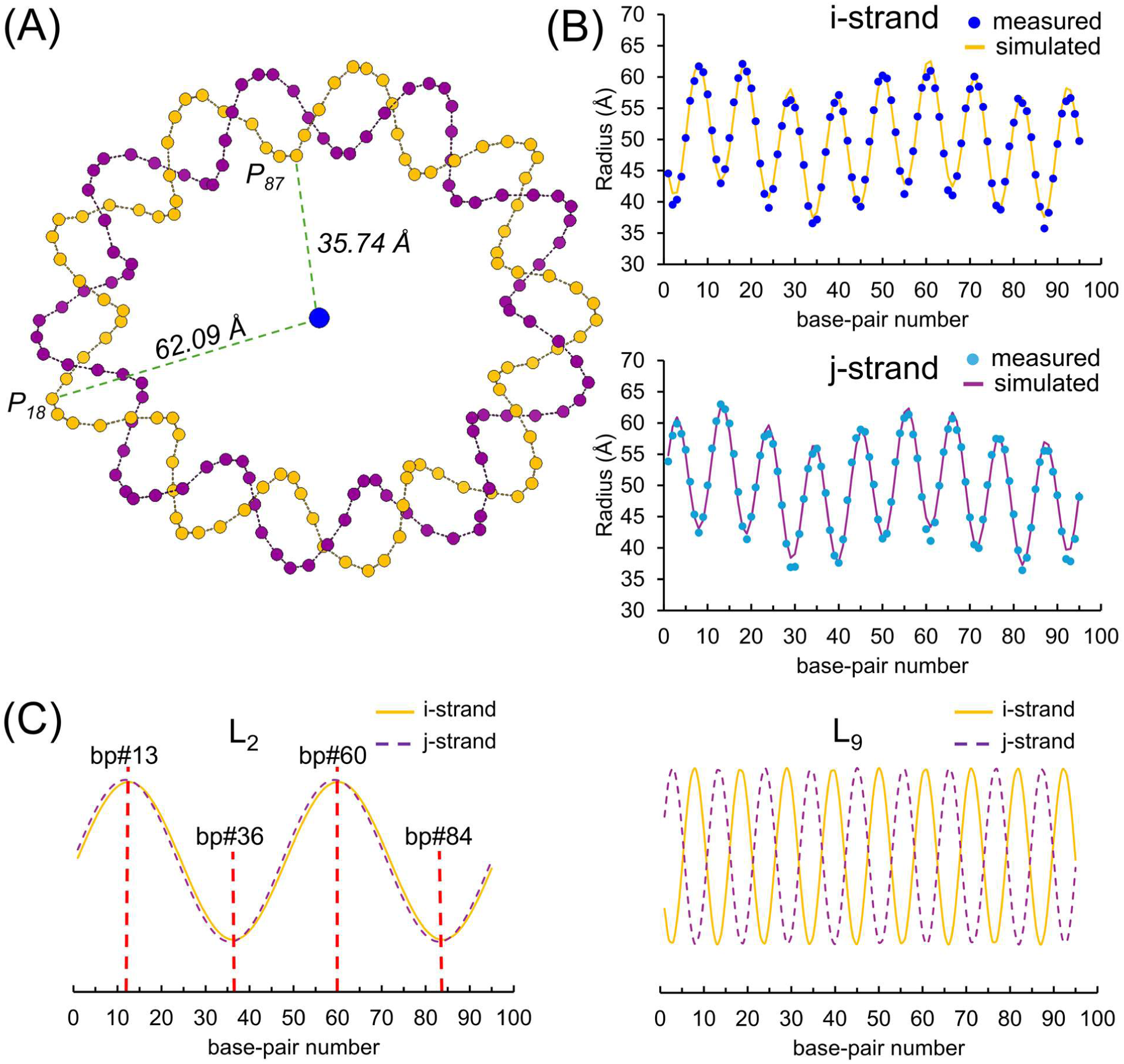
Analysis of helical twist of dsMC95. (A) Radii measurements. The phosphate atoms of the i-strand (orange dots) and the j-strand (purple dots) are plotted. The center-of-mass is represented by the blue dot. The dashed lines mark the shortest and longest radius, respectively, of the i-strand. (B) Radii vs. bas-pair-number plots for the i-strand (top) and the j-strand (bottom). “Simulated” lines were generated according to Eq. 1, and the respective parameters are: i-strand, a_0_=50.03Å, c_9_=-0.194±0.006, θ_1_=(2.2±1.8)°, c_2_=0.060±0.006, and θ_2_=(−5.5±5.8)°; j-strand, a_0_=50.04Å, c_9_=0.194±0.006, θ_1_=(−7.5±1.8)°, c_2_=0.062±0.006, and θ_2_=(0.4±5.6)°. (C) Overlay of the respective L_9_ (left) and L_2_ components. Maxima and minima of the L_2_ components are marked. See additional information in SI sect. S3.1.

Furthermore, major and minor grooves were analyzed following an established procedure (70). Major groove widths (majGW) at each base-pair of dsMC95 were computed by calculating the P-P distances between the appropriate pairs of P-atoms (Figure 5A, SI sect. S3.2). The resulting majGW values oscillate along the DNA ring with a variation pattern closely matching that of the distances (R_major_) between the mid-points of the corresponding PP pairs and the center-of-mass of the ring (Figure 5B). This indicates that outward-facing major grooves (i.e., large R_major_) are wider, while those inward-facing (i.e., small R_major_) are narrower. A similar oscillatory pattern is observed for minor groove widths (minGW, Figure 5C). Consistent with a B-DNA configuration of dsMC95, majGW are larger than minGW (Figure 5). Variations in the major and minor groove widths reflect bending of the DNA duplex imposed by circular confinement. Although this behavior has been proposed previously based on computation models of free DNA rings (55) as well as been observed in other bent-DNAs within protein-DNA complexes (71, 72), the analysis of dsMC95 provides the first direct assessment based on an experimentally determined structure of a DNA-only ring.

**Figure 5:**
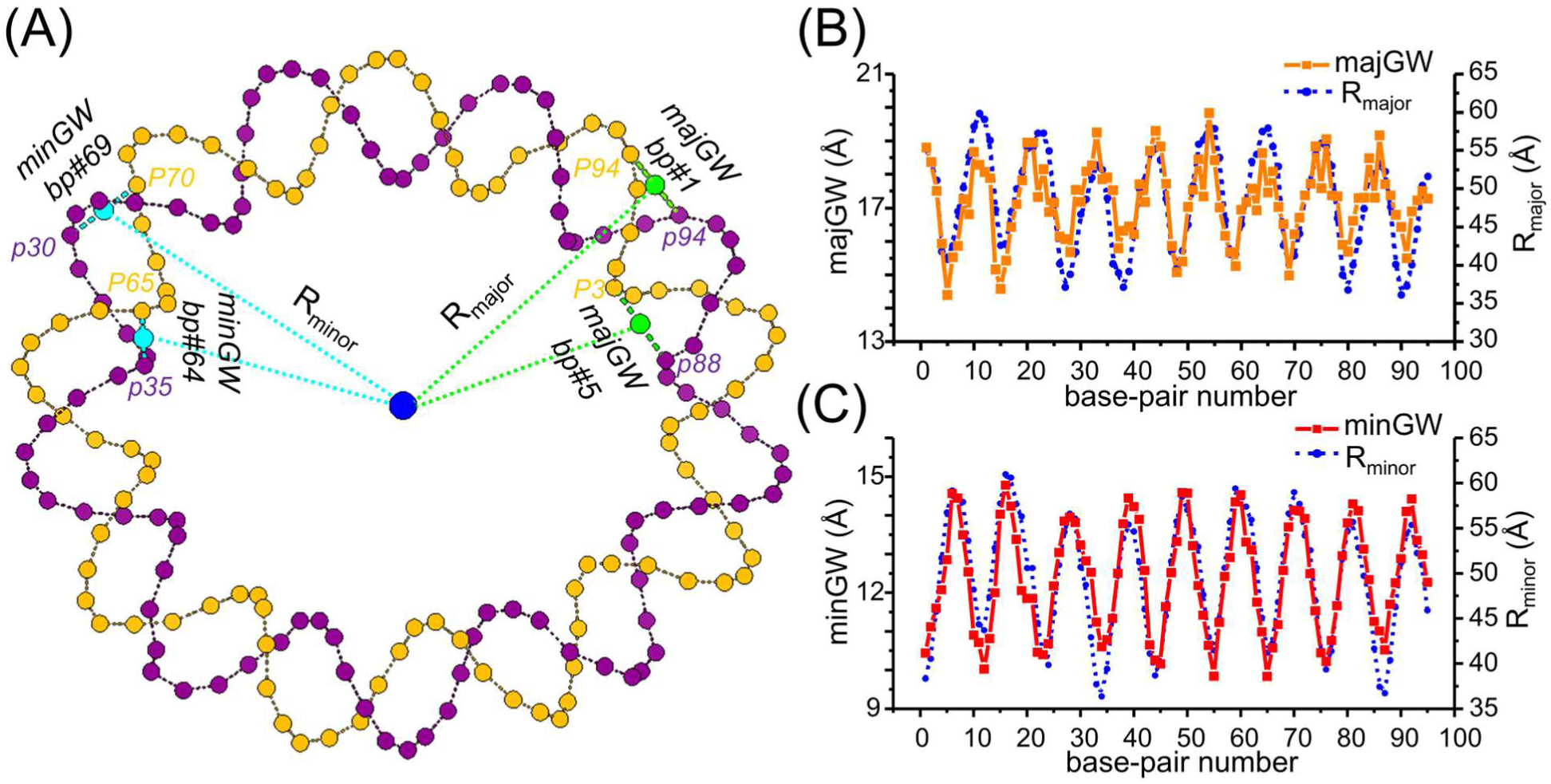
Analysis of groove widths. (A) Examples of groove width calculation. The phosphate atoms of the i-strand (orange dots) and the j-strand (purple dots) are plotted. The center-of-mass is represented by the blue dot. Examples shown for major grooves (green) are bp#1, majGW 18.8 Å, R_major_ 55.3 Å; and bp#5, majGW 14.4 Å, R_major_ 40.8 Å. Examples shown for minor grooves (blue) are bp#64, minGW 9.8 Å, R_minor_ 41.2 Å; and bp#69, minGW 14.1 Å, R_minor_ 59.0 Å. (B) Overlay of the majGW vs. bp# plot (orange) with the R_major_ vs. bp# plot (blue). (C) Overlay of the minGW vs. bp# plot (red) with the R_major_ vs. bp# plot (blue).

### Deformation of dsMC95 ring

To characterize deformation of the DNA ring, we calculated the mid-points of the P atom pairs associated with each base-pair (designated as mP, Methods; SI sect. S3.3, Figure S9A), which serve as proxies for the center of each base-pair. The mP points were transformed into an “internal” dsMC coordinate system (see Methods), with the origin set at the Mc of the DNA circle. Base-pair vector 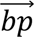(13) (i.e., 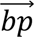(a), eq. 5), which corresponds to the base-pair at the first maximum in the L_2_ periodicity of the radii (Figure 4C), was set as the X-axis (i.e., 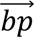(a), eq. 5). The second vector to define the XY-plane was chosen as 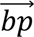(36) (i.e., vector 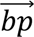(b), eq. 6), which corresponds to the base-pair at the first minimum of the L_2_ periodicity (Figure 4C). Upon establishing the orthogonal *̂x*, *̂y*, and *̂z* axes, transformations were carried out to obtain the dsMC coordinates for each mP (eq.s 8-9).

The in-plane shape of the dsMC95 ring, which is characterized by the X and Y coordinates of mP in the dsMC coordinate system, can be fitted well by an ellipsoid with a long-axis of 52.5±0.4 Å and a short-axis of 46.4±0.4 Å (Figure 6A). The computed ellipticity is 1.13±0.02, which is consistent with that measured directly from the cryo-EM map (Figure 3A, top-view, width/height ∼ 1.10). In addition, the long-axis (bp#13 as the +x axis and bp#60 as the -x axis) and the short-axis (approximately bp#36 as the +y axis and bp#84 as the -y axis) of the ellipsoid correspond, respectively, to the maximum and minimum of the L_2_ periodicity of the radii (Figure 4C). This indicates that the L_2_ periodicity arises primarily from the deviation of the DNA ring from a perfect circle.

**Figure 6:**
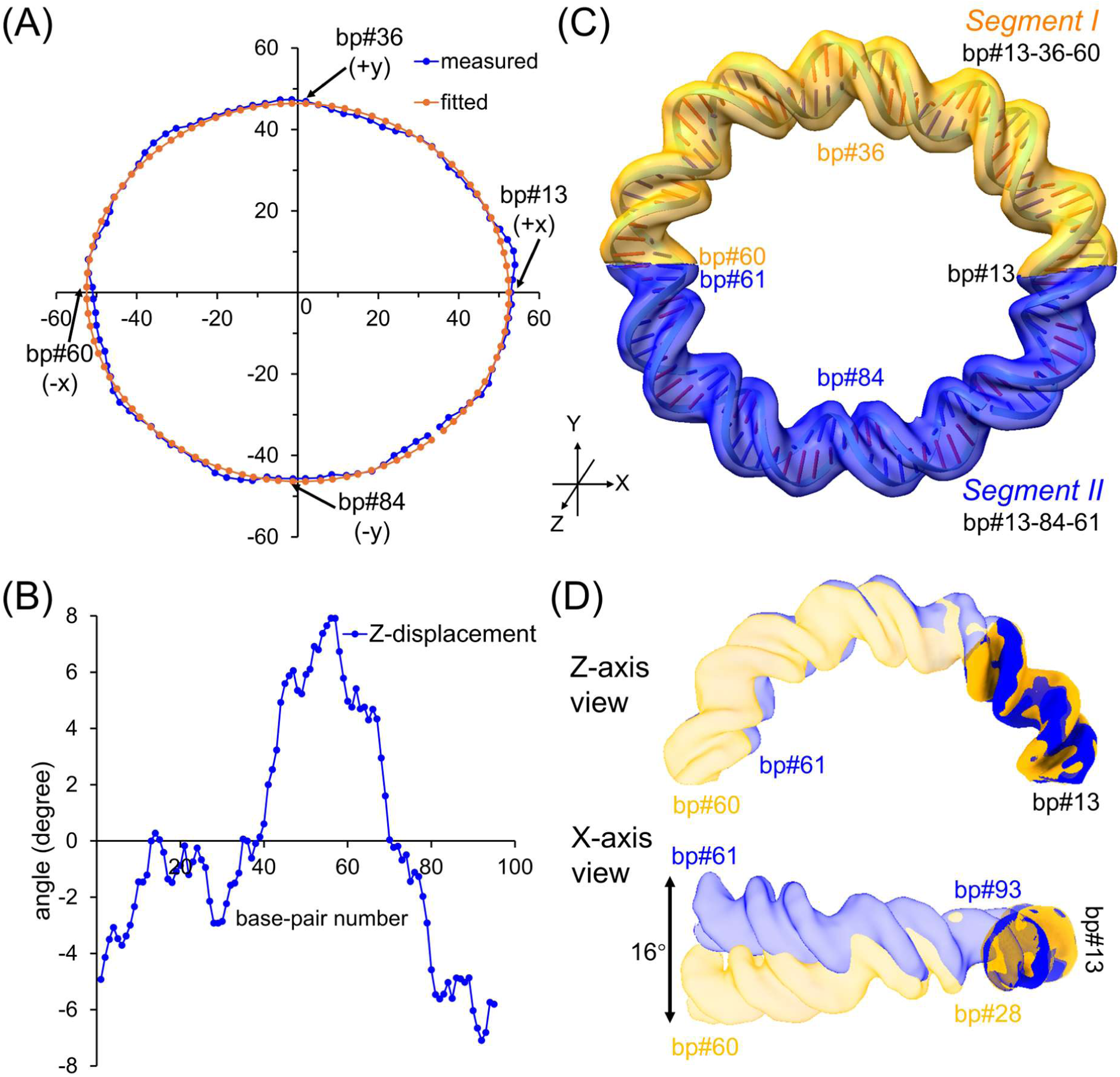
Analysis of dsMC95 deformation. (A) In-plane analysis. The blue dots represent the projections of base-pair midpoints (mP) onto the XY-plane in the internal dsMC coordinate system, and the orange line represents a fit to an ellipsoid (eq. 7, Methods). Parameters obtained from fitting are a= (52.5±0.4) Å, b = (45.4±0.4) Å, and φ_1_= (2.0±2.8)°. Base-pair numbers of mP located at the axes of the ellipsoid are marked. (B) Out-of-plane analysis. The mP displacement angles are computed according to eq. 5 (Methods) and plotted vs. the base-pair numbers. (C) Segmentation of dsMC95 map, with Segment I (bp#13-28-60) shown in orange and Segment II (bp#13-93-61) shown in blue. See additional information in SI, sect. S3.4. (D) Comparison of the two half-circle segments, with views along the Z-axis (top) and along the X-axis (bottom). Rigid body alignment was carried out based on alignment between the bp#13-28 sub-fragment of Segment I and the bp#13-93 sub-fragment of Segment II. CC between the two sub-fragments is 0.99.

The Z-coordinates of mP in the dsMC coordinate system represent displacements of the respective base-pair centers away from the XY-plane (SI sect. S3.3, Figure S9B). For the 95 mP points, the average displacement angle is -0.10° (Figure 6B), indicating that the chosen XY-plane nearly evenly cuts across the DNA ring (SI sect. S3.3, Figure S9B). The maximum and minimum tilt angles are +7.9° and -7.1° (Figure 6B, Figure S9B), respectively. This corresponds to an overall out-of-plane displacement of 15°, which is consistent with the predominantly flat ring observed in the cryo-EM map (Figure 3A, side-views).

While the mP approach described above used a reduced dimension to analyze the dsMC95 ring, the key features obtained are supported by direct analysis of the 5.3 Å cryo-EM map. Based on analysis in the XY-plane, the dsMC95 map was dissected into two segments, with Segment I assigned as bp#13-36-60 and encompassing the (+x)-(+y)-(-x) half, and Segment II assigned as #13-84-61 and encompassing the (+x)-(-y)-(-x) half (Figure 6C). The two segments are similar with an overall correlation coefficient (CC) of 0.97 (SI sect. S3.4, Figure S10). Nevertheless, when the two half-circle segments are overlaid and aligned at their bp#13 ends, their opposite ends diverge (Figure 6D). Specifically, when viewed along the Z-axis, only minimal differences are observed, indicating that bending within the XY-plane is nearly identical between the two halves (Figure 6D, top). In contrast, when viewed along the X-axis, the two segments tilt away from each other with deviations becoming apparent approximately 15 base-pairs away from the aligned ends (i.e., bp#28 in Segment I and bp#93 in Segment II) (Figure 6D). The displacement angle measured directly from the map is approximately 16° (Figure 6D), which is in close agreement with the value of 15° obtained from the mP analysis (Figure 6B). Overall, the analyses indicate that deformations of dsMC95 are modest, characterized by slight ellipticity and minor out-of-plane displacement.

### Comparison between dsMC95 and other DNAs

As an attempt to gain understanding of physical features of DNA duplexes such as bending, the dsMC95 cryo-EM map was compared with other DNA structures. For example, the Feigon lab has solved NMR structures of two 12 base pair DNA duplexes formed by d(GCAAAATTTTGC) ([A4T4], PDB ID 1RVH) and d(CGTTTTAAAACG) ([T4A4], PDB ID 1RVI) (73). Due to the difference at the central ApT vs. TpA step, the [A4T4] DNA shows an overall bend, while the [T4A4] DNA is straight (73). Using ChimeraX, density maps at 5 Å-resolution for each of these structures were generated and then aligned with that of dsMC95 (SI sect. S4.1). Aligning the [T4A4] DNA with dsMC95 yields a maximal CC of 0.85, with some deviations observed between a helical turn of dsMC95 and the terminal regions of [T4A4] DNA (Figure 7A, SI sect. S4.1, Figure S13). This indicates differences in DNA curvature between the dsMC95, which bends to form the closed circle, and the [T4A4] duplex, which is overall straight. On the other hand, the [A4T4] DNA can be aligned with dsMC95 with a maximal CC of 0.91 (Figure 7A, SI sect. S4.1, Figure S14), indicating that this bent DNA conformation fits better with a helical turn within the dsMC95 ring.

**Figure 7:**
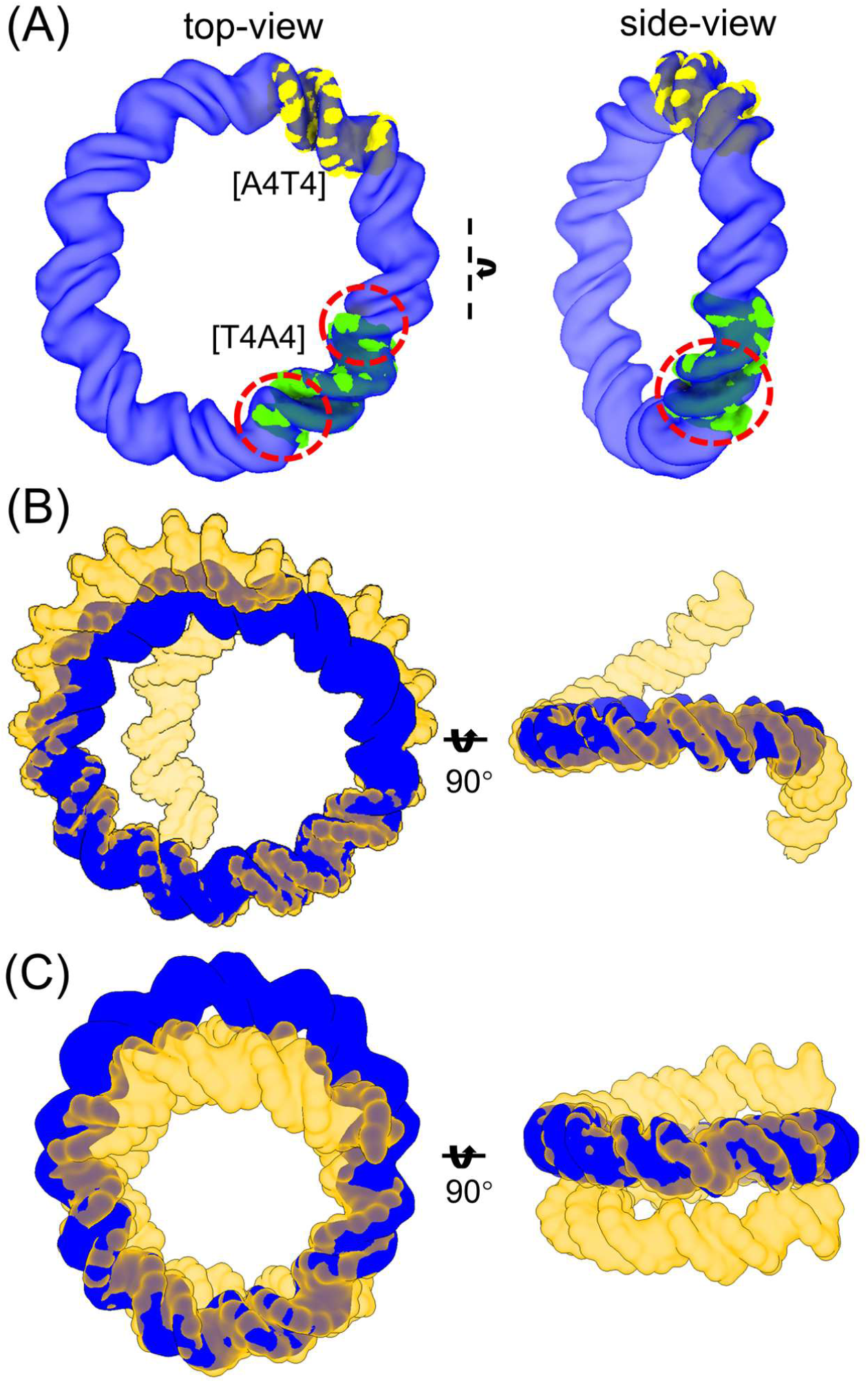
Comparison of dsMC95 to other DNAs. In each panel, the top-view is shown on the left, and a side-view is shown on the right. (A) Optimal overlay between the dsMC95 map (blue) and the 5Å maps rendered from NMR structures of [T4A4] (1RVI.pdb, green) and [A4T4] (1RVH.pdb, yellow). The dash circles mark the most severe differences between dsMC95 and [T4A4]. See additional information in SI sect. S4.1. (B) Overlay between the dsMC95 map (blue) and the 5Å map rendered from a model of a d(CA4T4G)_15_ duplex containing 15-repeat of [A4T4] (yellow). The side-view of d(CA4T4G)_15_ shows only a portion of the map. See additional information in SI sect. S4.1. (C) Optimal overlay between the dsMC95 map (blue) and the 5Å map rendered from the nucleosomal DNA structure presented 1KX5.pdb (orange). See additional information in SI sect. S4.2.1. For Table of Contents Only

Furthermore, NMR structure-based modeling has shown that DNA with repeating [A4T4] units forms a superhelix consisting of ∼ 120 bp/turn (73). We therefore generated a model of a duplex formed by d(CAAAATTTTG)_15_ ([A4T4]_15_, i.e., 15-repeating [A4T4] units) and rendered it into a 5Å map (SI sect. S4.1) for comparison with dsMC95. The overlay shows that the [A4T4]_15_ model aligns well with a 45-base-pair segment of dsMC95 (i.e., bp#14-bp#59) that corresponds to nearly half of the ring (Figure 7B). The in-plane diameter of dsMC95 is approximately 90% of that of [A4T4]_15_ (Figure 7B), indicating a similar degree of DNA bending. On the other hand, more pronounced differences can be observed along the Z-axis (Figure 7B), where the unconstrained [A4T4]_15_ adopts a super-helical conformation with a large out-of-plane span, in contrast to the predominantly planar dsMC95 (Figure 7B).

Comparisons with free DNAs are limited because atomic-resolution structures have been experimentally determined only for linear duplexes shorter than two helical turns. We therefore compared our map with longer DNA duplexes observed in structures of protein-bound complexes. One of the examples is the nucleosomal DNA, which wraps around the histone core to form the nucleosome core particle. The atomic structure of a 147-base-pair DNA duplex was isolated from a crystal structure of the canonical nucleosome core particle, PDB ID 1KX5 (74), which has been used as a well-established structure for studying DNA bending within the nucleosome. The 1KX5 DNA was rendered into a 5Å density map using ChimeraX (SI sect. S4.2.1) and compared with the dsMC95 map (Figure 7C). The two maps superimpose well over a 34-base-pair segment, with a CC of 0.96 and closely aligned major and minor grooves (Figure 7C; SI sect. S4.2.1 Figure S15B). This indicates similar DNA bending between dsMC95 and the 1KX5 DNA over approximately three helical turns (or about one-third of the dsMC95 ring). Notable differences emerge when comparing the entire dsMC95 map with that of the 1KX5 nucleosomal DNA. The 1KX5 nucleosomal DNA shows more pronounced overall bending as its in-plane diameter is approximately 10% smaller than that of dsMC95 (Figure 7C, top-view). This increased curvature is partly accommodated by a greater span of the nucleosomal DNA along the Z-axis (Figure 7C, side-view) as well as by interactions with the histone core. Similar results were obtained when compared to another structure of nucleosomal DNA (SI, sect S4.2.2).

Overall, these comparisons of dsMC95 with other DNA structures indicate that the bending required to form the dsMC95 ring falls within the range observed for unconstrained linear duplexes and may be further modulated by the covalently connected termini or protein-DNA interactions.

## Discussion

We report here a 95-bp circular DNA structure captured at a resolution of 5.3 Å (Figure 3, SI sect. S2), which currently represents the only cryo-EM structure of a free DNA with a resolution better than 6 Å. dsMC95 forms a closed-ring double-stranded DNA structure with nine helical turns of B-form duplex, with no observable local deformations (Figures 3, 4, and 5). The DNA ring exhibits a predominantly circular and planar conformation (Figure 3), with an in-plane ellipticity of approximately 1.13 and an out-of-plane displacement of approximately 15° (Figure 6). Comparison of dsMC95 with other DNA structures revealed both similarities and differences in DNA bending (Figure 7). The results advance our understanding of DNA structure under topological constraints.

Previously, small DNA circles have been extensively used to investigate physical properties of DNA duplexes. Most relevant to this work, Demurtas and co-workers have investigated bending of short DNA duplexes by carrying out cryo-EM studies of 94-bp DNA minicircles that are either covalently closed, doubly nicked, or doubly gapped (38). The reconstructed micrographs, although limited by their nanometer resolutions, revealed that the DNA rings were planar, and the covalently-closed DNAs showed no kinks (38). Our findings are consistent with these observations (Figure 3 and Figure 6). Interestingly, in Demurtas’ work the average ellipticity for the covalently-closed circles was reported to be 1.16±0.09 and 1.22±0.13, respectively, with and without Mg^2+^ (38). This is very similar to the ellipticity of 1.13±0.02 determined for dsMC95 (Figure 6A). Note that our dsMC95 structure was obtained in the presence of Tris-EDTA buffer without additional salt (see Methods). As previously reported, in such a “low salt” environment, the negatively charged phosphate groups along the DNA backbone are not effectively screened, and their resulting electrostatic repulsion promotes a relatively symmetric circular and planar shape of these small DNA circles (38, 49). The strong repulsion may also affect the local environment of the base pairs as revealed from the 2AP fluorescence measurements (Figure 2B).

The improved resolution of dsMC95 provides information on the DNA helix under the constraints imposed by the covalently-closed ring. The map shows both strands of the DNA with clearly distinguishable major and minor grooves (Figure 3). Analysis of the DNA backbone using coordinates of the phosphate atoms (Figures 4, 5, 6) reveals a number of features. First, the phosphate atoms show a clear periodicity of nine in the helical twist of each DNA strand that is out-of-phase between the two strands (Figure 4B and 4C, SI sect. S3.1). This has been previously proposed based on analysis of computer-generated models (55, 57), but could not be assessed with low-resolution EM maps (38, 48, 49). Furthermore, the resulting 10.56-bp per turn in helical twist indicates a B-DNA duplex, which is consistent with conclusions drawn previously from modeling and DNA cyclization measurements (35). Second, groove widths of dsMC95 are found to vary periodically, narrowing when facing the center of the ring and widening when oriented outward (Figure 5, SI sect. S3.2). This has been described previously in computer modeling studies (55), but has not been observed experimentally in free DNA. Because variations in DNA curvature (i.e., the inverse of radius) and groove width significantly impact functions such as protein recognition (28, 55), the ability to experimentally determine these features is highly desirable. In addition, by developing a new algorithm to trace the DNA helix using the mid-points of phosphate atoms within each base-pair (Figure 6, SI sect. S3.3), we were able to identify local DNA variations, including both in-plane ellipticity (Figure 6A) and out-of-plane displacement (Figure 6B). Indeed, the dsMC95 map provides sufficient resolution to reveal small but measurable variations between the two half-segments of the DNA ring (Figure 6C and 6D).

The dsMC95 map allowed comparison of the minicircle with other DNA structures (Figure 7). The intrinsically bent [A4T4] DNA aligned with dsMC95 better than the straight [T4A4] DNA (Figure 7A, SI sect. S4.1). This reinforces the notion that bending required for forming the dsMC95 circle is within the realm of unconstrained duplexes (35, 38). It also raises an intriguing question regarding how DNA bending is shaped by the interplay between intrinsic sequence-specific properties and topological constraints. In addition, DNA minicircles have been proposed to mimic DNA wrapped around histones in nucleosome core particles (28, 49). Comparisons between dsMC95 and representative nucleosomal DNA structures provide the first experimental assessment (Figure 7C, SI sect. S4.2). The analysis reveals similarity in bending between dsMC95 segments and nucleosomal DNAs. However, the overall in-plane diameter of the nucleosomal DNA is smaller than that of dsMC95, indicating tighter curvature that is accommodated by out-of-plane displacement, which is possible partly because the nucleosomal DNA has unconstrained termini. These findings highlight the role of topology, in addition to protein-DNA interactions, in modulating DNA duplex conformations such as bending.

While the dsMC95 map resolution reported in this work is substantially improved from prior EM studies of DNA circles (11, 12, 38, 48–51) and surpasses all DNA-only structures currently deposited in EMBD (58–60), it is not yet sufficient for assigning the individual bases. dsMC95 lacks distinct local feature(s) that could facility cryo-EM reconstruction, and attempts to improve resolution by symmetry averaging (data not shown) were not successful, which likely is due to the lack of symmetry in the DNA sequence used. In other reported free DNA cryo-EM studies, the presence of termini allowed assignment of the entire sequence into maps with resolutions worse than 7 Å (60). However, the same approach is not applicable to dsMC95 due to the circular nature of this DNA. Further improvements in map resolution that enable assignment of individual bases would facilitate a deeper understanding of sequence-specific features of DNA under torsional constraints. Accordingly, continued methodological advances are highly desirable.

In summary, the dsMC95 structure reported in this work represents the first cryo-EM study of a free DNA with a resolution better than 6 Å. The analysis shows that the global conformation of the DNA minicircle is consistent with computational models and previous studies of closed circular DNAs. It further reveals structural details of the duplex that advance our understanding of DNA structure under topological constraints. The knowledge gained should benefit efforts on understanding and manipulating circular DNA in biological systems as well as in nanotechnology developments.

## Acknowledgements

Electron microscopy data were collected at the Core Center of Excellence in Nano Imaging (CNI) at University of Southern California (USC). Cryo-EM data processing was carried out at the Center for Advanced Research Computing (CARC) at USC. We thank Dr. H. Khant (USC) for assisting with cryo-EM data acquisition, and Dr. Tomek Osinski (USC) for assisting with cryo-EM data processing, and Dr. J. Jiang (NIH/NHLBI) for simulating discussions on analysis of cryo-EM data and dsMC95 structural features.

## Author Contributions

Y.L. and P.Z.Q. designed the studies and wrote the manuscripts; Y.L. designed, synthesized, biochemically characterized dsMC95, and carried out cryo-EM data acquisition and analysis. H.C. assisted in biochemical analysis of dsMC95. Y.H. and J.F. assisted in cryo-EM data analysis. Y.L., K.L., D.K., and V.C. contributed to model building. Y.L., P.Z.Q, and J.F. carried out analysis of dsMC95 models and comparison with other DNAs. All authors reviewed the manuscript.

## Supplementary Data

Supplementary Data is available online.

## Data Availability

The atomic model and the cryo-EM map have been deposited and will be released upon publication of the peer-reviewed article following wwPDB policies.

---

For Table of Contents Only

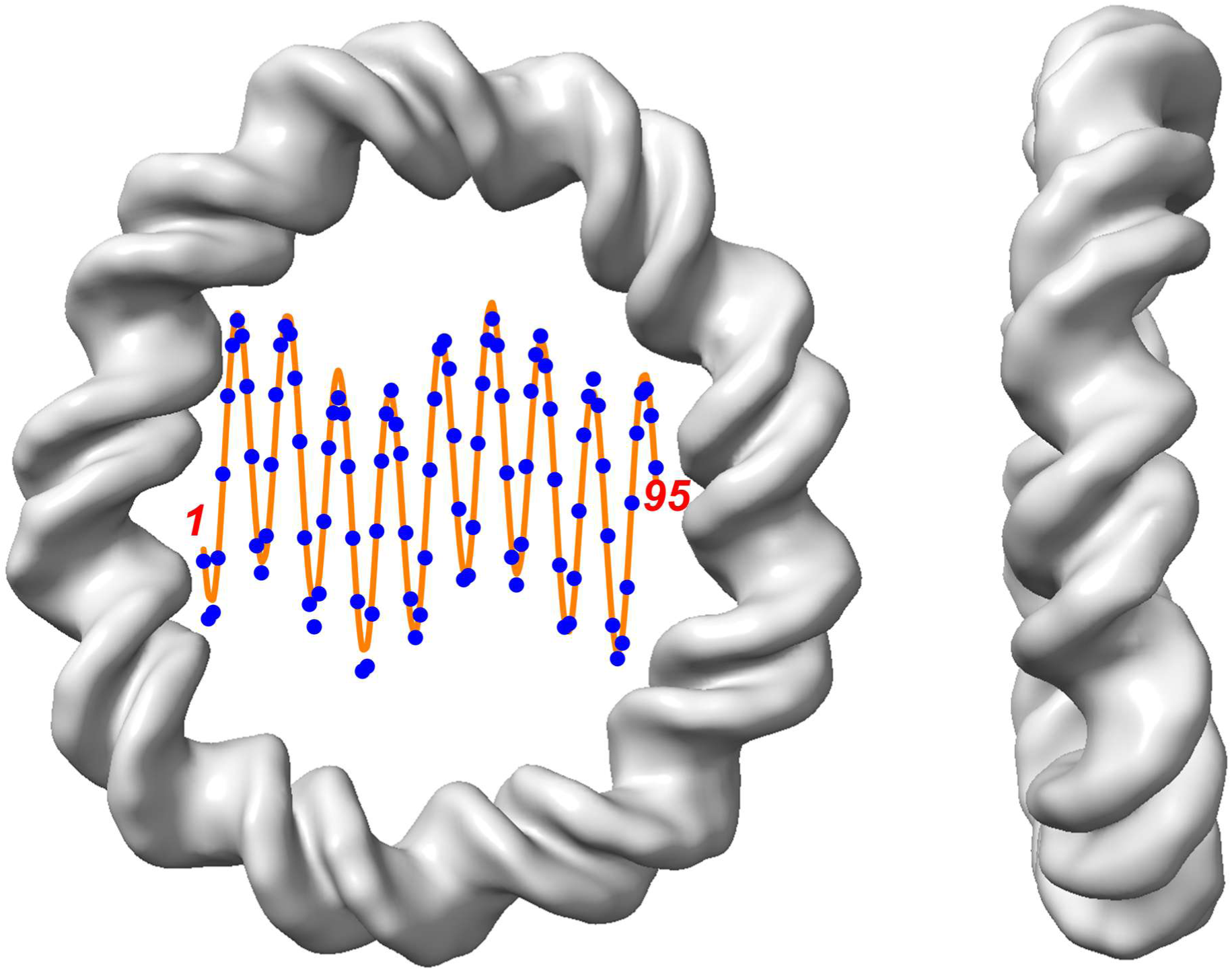

