## Supplemental Information for "Cryo-EM Structure of a 95-Basepair Double-Stranded DNA Minicircle at 5.3 Å Resolution"

### Cryo-EM Structure of a 95-Base-Pair Double-Stranded DNA Minicircle at 5.3 Å Resolution

Table of Contents

**S1. Additional information on dsMC95 synthesis and characterization**
- S1.1. Additional information on design of dsMC95
- S1.2. Additional biochemical characterization of dsMC95
- S1.3. Additional information on 2-amino-purine fluorescence measurement

**S2. Additional information on cryo-EM data collection and analysis**

**S3. Additional information on analysis of dsMC95 conformation**
- S3.1. Analysis of radius
- S3.2. Analysis of groove width
- S3.3. Analysis using the dsMC coordinate system
- S3.4. Variations observed directly from cryo-EM map

**S4. Additional information on comparison between dsMC95 and other DNA**
- S4.1. Comparison of dsMC95 with other free DNA
- S4.2. Comparison of dsMC95 with nucleosomal DNA

#### S1. Additional information on dsMC95 design and characterization

##### *S1.1. Additional information on design of dsMC95*

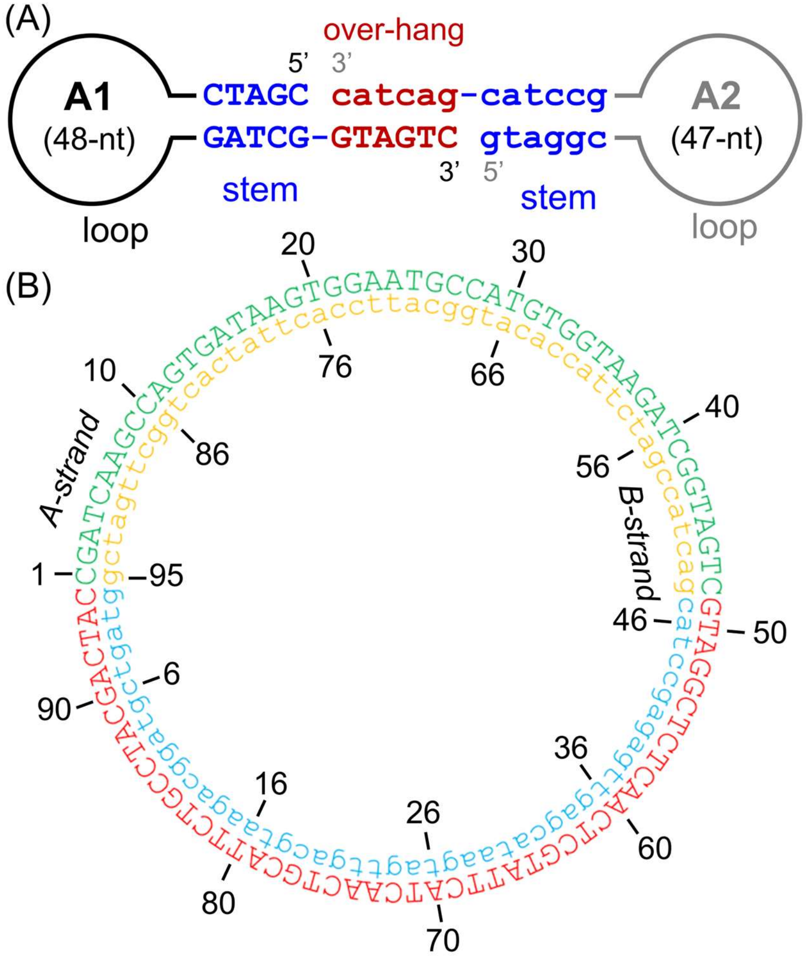

**Figure S1:** Design of dsMC95. (A) A schematic representation of the A1 and A2 hairpins used to generate a single-stranded minicircle (ssMC). The stem (blue) and overhang (red) sequences of A1 and A2 are shown in upper and lower cases, respectively. The overhang segments hybridize to form the intermediate nicked DNA dumbbell (see Step 1 in Figure 1, main text). Installation of a 5'-phosphate at both strands allows ligation to seal the two 5'-3' gaps to generate a ssMC. The same scheme is also applicable for the complementary B1 and B2 strands. Detailed sequences are listed in Table S1. (B) A complete primary sequence of dsMC95. The A-strand sequence is shown in uppercase letters (outer circle), with nucleotide numbering increasing in the 5′→3′ direction. This strand is formed by ligation of A1 and A2 and is also referred to as A-ssMC in its single-stranded form. The complementary B-strand sequence is shown in lowercase letters, with nucleotide numbering increasing in the 5′→3′ direction. This strand is formed by ligation of B1 and B2 and is referred to as B-ssMC in its single-stranded form. Color coding marks the respective sequence of each strand: A1 (green), A2 (red), B1 (orange), and B2 (blue). Base-pair numbering is defined based on the sequence of the A1 strand. Specifically, base pair #1 corresponds to pairing between C1 of the A-strand (nucleotide at the 5'-terminus of the A1 strand, Table S1) and g95 of the B-strand (nucleotide at the 3'terminus of the B1 strand, Table S1).

Following the scheme presented in Figure 1 of the main text, the 95 base-pair (bp) double-stranded minicircle (dsMC95) was synthesized using four oligonucleotides designated as A1, A2, B1, and B2. Figure S1A shows the key nicked dumbbell intermediate formed by A1 and A2, and Figure S1B shows the complete primary sequence of dsMC95. Detailed sequences of the DNA strands are listed in Table S1.

**Table S1.** Sequence of individual DNA strands

| Name | Sequence (5' – 3') |
|---|---|
| A1 (48-nt)[(a)] | CGATCAAGCCAGTGATAAGTGGAATGCCATGTGGTAAGA<br>TCGGTAGTC |
| A2 (47-nt) | GTAGGCTCTCAACTCGTATTCATCAACTGCATTCTGCCTA<br>CGACTAC |
| B1 (48-nt) | GACTACCGATCTTACCACATGGCATTCCACTTATCACTGG<br>CTTGATCG |
| B2 (47-nt) | GTAGTCGTAGGCAGAATGCAGTTGATGAATACGAGTTGA<br>GAGCCTAC |
| A1-40-FAM[(b)] | CGATCAAGCCAGTGATAAGTGGAATGCCATGTGGTAAGA/<br>**FAM_dT/**CGGTAGTC |
| A1-18-2AP[(c)] | CGATCAAGCCAGTGATA/**2AP**/GTGGAATGCCATGTGGTAA<br>GATCGGTAGTC |
| A-linear (95-nt)[(d)] | CGATCAAGCCAGTGATAAGTGGAATGCCATGTGGTAAGA<br>TCGGTAGTCGTAGGCTCTCAACTCGTATTCATCAACTGCA<br>TTCTGCCTACGACTAC |
| A'-linear (95-nt)[(e)] | GTAGGCTCTCAACTCGTATTCATCAACTGCATTCTGCCTA<br>CGACTACCGATCAAGCCAGTGATAAGTGGAATGCCATGT<br>GGTAAGATCGGTAGTC |
| B-linear (95-nt)[(f)] | GTAGTCGTAGGCAGAATGCAGTTGATGAATACGAGTTGA<br>GAGCCTACGACTACCGATCTTACCACATGGCATTCCACTT<br>ATCACTGGCTTGATCG |
| B'-linear (95-nt)[(g)] | GACTACCGATCTTACCACATGGCATTCCACTTATCACTGG<br>CTTGATCGGTAGTCGTAGGCAGAATGCAGTTGATGAATAC<br>GAGTTGAGAGCCTAC |

(a) The nucleotide at the 5'-terminus of the A1 strand was set as nucleotide #1 of the A-strand in dsMC95 (Figure S1B).
(b) A fluorescent dT (6-FAM) substitution is indicated by the bolded "**/FAM_dT/**". Within dsMC95, the fluorescence label is located at nucleotide #40 of the A-ssMC (Figure S1B).
(c) A 2-aminopurine substitution is indicated by the bolded "**/2AP/**". Within dsMC95, the fluorescence label is located at nucleotide #18 of the A-ssMC (Figure S1B). This is designated as site-1 as described in Figure 2B of the main text.
(d) 95-nucleotide (nt) linear single-stranded DNA generated by ligating an A1 strand with a 5'-OH group with an A2 strand with a 5'-phosphate group. The sequence of A1 is underlined.
(e) 95-nt linear single-stranded DNA generated by ligating an A1 strand with a 5'-phosphate group with an A2 strand with a 5'-OH group. The sequence of A1 is underlined.
(f) 95-nt linear single-stranded DNA generated by ligating a B1 strand with a 5'-phosphate group with a B2 strand with a 5'-OH group. The sequence of B1 is underlined.
(g) 95-nt linear single-stranded DNA generated by ligating a B1 strand with a 5'-OH group with a B2 strand with a 5'-phosphate group. The sequence of B1 is underlined.

***S1.2. Additional biochemical characterization of dsMC95***

*S1.2.1. Characterization of yields in DNA circle synthesis reactions*

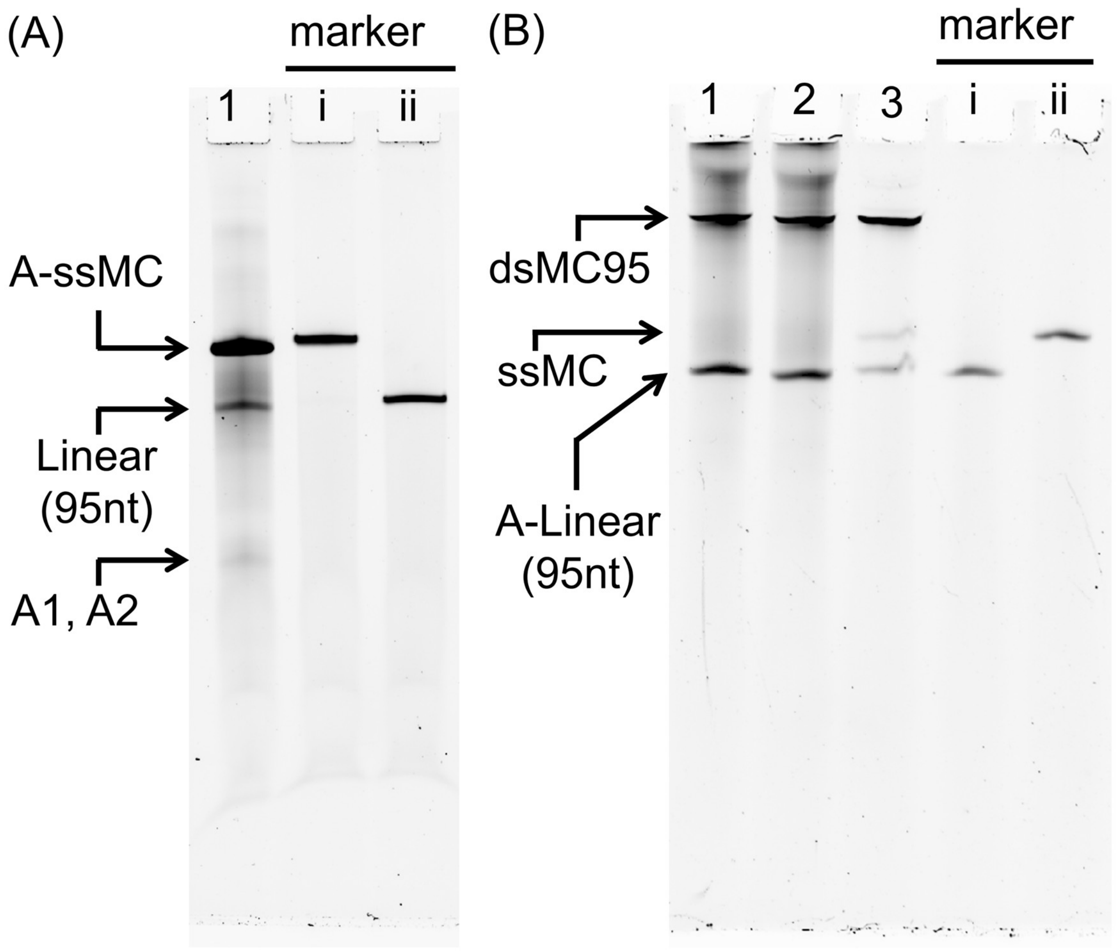

**Figure S2**: Characterization of DNA circle synthesis. DNA samples were resolved on a 10% polyacrylamide denaturing gel, and were visualized by SYBR Gold staining. (A) Characterization of ssMC synthesis. Lane 1 shows crude reaction product from a synthesis of A-ssMC (see Figure S1B for sequence), in which the A1 strand containing a 5-phosphate group was ligated with the A2 strand containing a 5'-phosphate group as described in Methods in the main text. The byproduct 95-nt linear DNA may include a mixture of A-linear and A'-linear DNAs (Table S1). The marker lanes are: (i) purified A-ssMC; (ii) purified A-linear. (B) Characterization of dsMC95 synthesis. Following Methods described in the main text, dsMC95 was synthesized from purified B-ssMC (see Figure S1B) and A-linear (Table S1) with various B-ssMC/A-linear ratios, and crude reaction products were characterized. Lane 1: B-ssMC/A-linear = 1/1.2; Lane 2: B-ssMC/A-linear = 1/1.1; Lane 3: B-ssMC/A-linear = 1/1. The marker lanes are: (i) purified A-linear; and (ii) purified A-ssMC. The percentages of dsMC95 among the total DNA are: Lane 1, 92%; Lane 2, 91%; and Lane 3, 87%. In lanes 1 and 2, bands above dsMC95 likely represent by-products.

Figure S2A shows an example of denaturing gel characterization of the crude product obtained in a synthesis of a single-stranded DNA circle. Besides the starting materials (A1 and A2, Figure S2A, lane 1) and the desired A-ssMC product, only a small amount of by-product was observed

(Figure S2A, lane 1). The reaction yield, defined as the percentage of the A-ssMC species among the total DNA, was 91% (Figure S2A, lane 1).

Figure S2B shows an example of denaturing gel characterization of the crude product obtained in a synthesis of dsMC95. Yields were above 85% with only small among of by-products (Figure S2B). Overall, the results show successful synthesis of DNA circles.

*S1.2.2 Chromatography characterization of dsMC95*

To further characterize DNA minicircles, a purified intact (fully covalently ligated) dsMC95, a nicked dsMC95, and a linear 95-bp DNA duplex of identical sequence were analyzed by size-exclusion chromatography (SEC). Figure S3 shows representative SEC traces for the three DNA constructs. Although all samples have identical sequence, the nicked dsMC95 eluted slightly earlier than the intact dsMC95, while the linear duplex exhibited a substantially longer retention time. These variations reflect different shapes of the three DNA in solution.

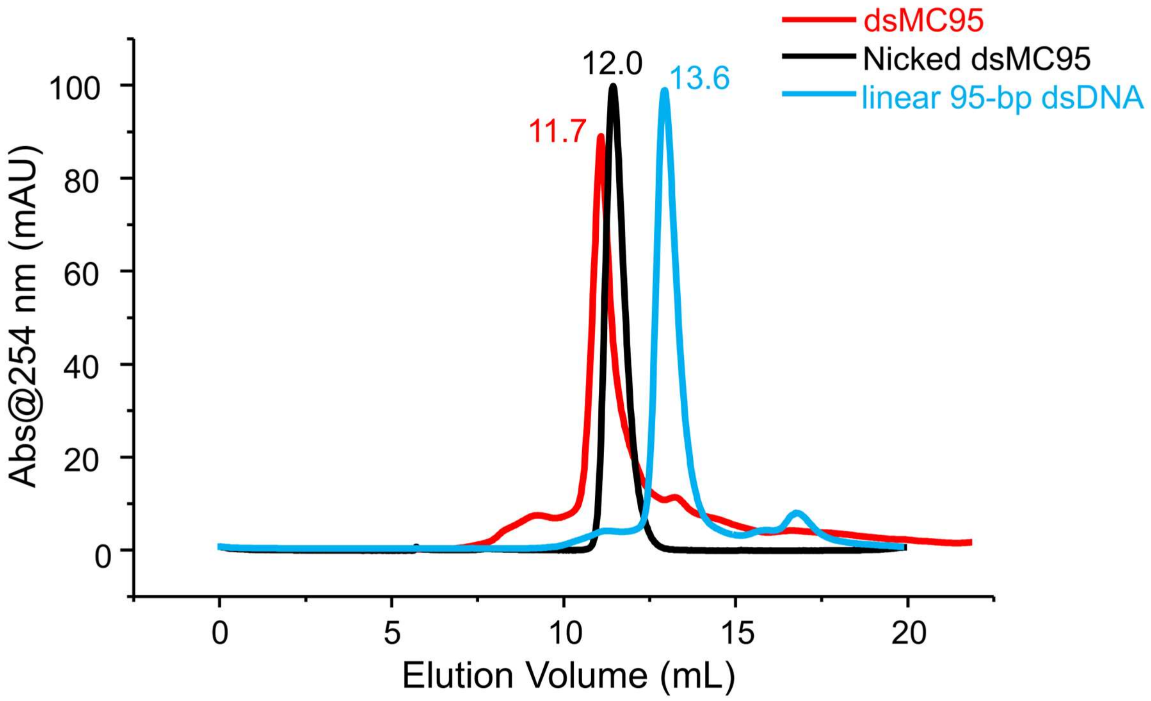

**Figure S3.** DNA circles analyzed by size-exclusion chromatography. Traces were acquired at 4 °C using an ÄKTA FPLC system equipped with a Superdex 200 Increase 10/300 GL column. The elution buffer was 20 mM Tris, pH 7.5 and 100 mM KCl, and the flow rate was 0.4 mL/min. The intact dsMC95 corresponds to the fully covalently ligated double-stranded minicircle (see Figure S1B). The nicked dsMC95 was generated by annealing B-ssMC (Figure S1B caption) with the A-linear strand (Table S1). The linear 95-bp duplex was prepared by annealing the A-linear and B-linear strands (Table S1).

##### *S1.3. Additional information on 2-aminopurine fluorescence measurement*

Following a previously established procedure (1), in this work, 2-aminopurine (2AP) fluorescence (F) and absorbance at 260 nm ($A_{260}$) were measured on the same sample:

$$F = k \cdot I_0 \cdot \varepsilon_{2AP} \cdot b \cdot c \cdot \varphi \qquad \text{(S1)}$$

$$A_{260} = \varepsilon \quad \cdot l \cdot c \qquad \text{(S2)}$$

where $k$ is the proportionality constant for the fluorimeter, $I_0$ is the incident light intensity, $b$ is the fluorescence path length, $c$ is the concentration of the sample, $\varepsilon_{2AP}$ is the extinction coefficient of the 2AP at the fluorescent excitation wavelength (320 nm in this work), φ is the quantum yield of the 2AP, ε is the extinction coefficient of the sample at 260 nm, and $l$ is the absorption path length.

The $F/A_{260}$ for a given sample is:

$$\frac{F}{A_{260}} = \frac{k \cdot I_0 \cdot \varepsilon_{2A} \quad \cdot b \cdot c \cdot \varphi}{\varepsilon \quad \cdot l \cdot c} = K \cdot \frac{\varepsilon_{2AP}}{\varepsilon} \cdot \varphi \qquad \text{(S3)}$$

where K is a proportional constant. As such, the $F/A_{260}$ value is proportional to the 2AP quantum yield ($\varphi$) but independent of the sample concentration:

**Table S2**: Additional information on $F_{370}/A_{260}$ measurements.[(a)]

| DNA construct | $F_{370}/A_{260}$ ($\times 10^4$) (a.u.) | | | |
|---|---|---|---|---|
| | Repeat 1 | Repeat 2 | Repeat 3 | Ave ± SD |
| dsMC95 | 0.43 | 0.17 | 0.55 | 0.40 ± 0.20 |
| Nicked MC95[(b)] | 3.8 | 4.2 | 4.1 | 4.0 ± 0.22 |
| Linear dsDNA[(c)] | 4.7 | 4.3 | 4.7 | 4.7 ± 0.39 |

(a) 2AP was incorporated at nucleotide #18 of the A-strand as shown in Figure 2B in the main text. Measurements carried out as described in Methods in the main text. "Ave ± SD" are reported in Figure 2B in the main text.
(b) a nicked MC generated by annealing B-ssMC (Figure S1B caption) and A-linear (Table S1).
(c) A linear double-stranded duplex generated by annealing A-linear and B-linear (Table S1).

Table S2 shows the individual measurements obtained for $F_{370}/A_{260}$, with $F_{370}$ being the fluorescence emission at 370 nm. The values were used to generate data presented in Figure 2B in the main text.

In addition, the $F/A_{260}$ approach was previously validated using a Fluorolog spectrofluorometer for fluorescence measurements and a LAMBDA UV/Vis/NIR spectrophotometer for absorbance measurements (1). In the present work, fluorescence measurements were carried out on a SpectraMax iD3 plate reader, while absorbance measurements were obtained using the same LAMBDA UV/Vis/NIR spectrophotometer. As shown in Figure S4, with the plate reader, control measurements gave the same $F/A_{260}$ values for DNA samples with different concentrations. This confirms the effectiveness of the $F/A_{260}$ approach for removing the dependence on sample concentration.

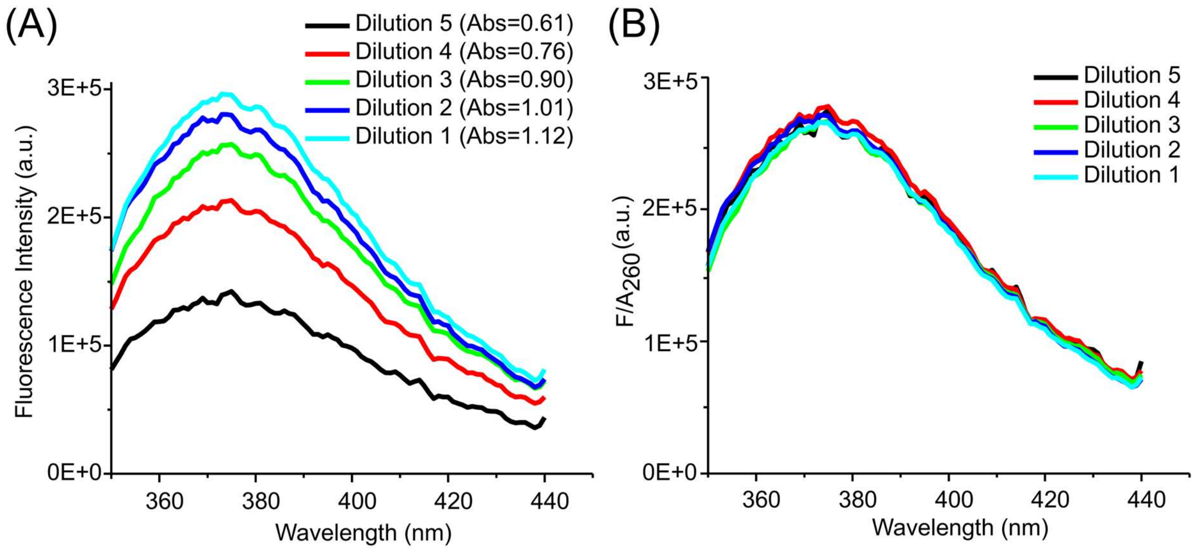

**Figure S4**. Evaluating the effectiveness of $F/A_{260}$. Data were obtained using the "A1-18-2AP" strand (Table S1) dissolved in a buffer containing 20 mM Tris, pH 7.5, 100 mM KCl. Fluorescence was measured at room temperature using a SpectraMax iD3 plate reader, and absorbance was obtained using a LAMBDA UV/Vis/NIR spectrophotometer (see Methods, main text). (A) Fluorescence emission spectra obtained for samples underwent serial dilutions from the same stock solution. The corresponding $A_{260}$ values for each sample are listed together with the sample number. (B) $F/A_{260}$ spectra derived from those shown in panel (A). These spectra overlay nicely, and the average $F_{370}/A_{260}$ value measured was $2.7 \times 10^5$, with a standard deviation of $0.1 \times 10^5$.

#### S2. Additional information on cryo-EM data collection and analysis

High-resolution cryo-EM data were acquired on a Titan Krios operating at 300 keV, yielding over 30,000 micrographs (see Methods, main text). Figure S5 summarizes the data processing workflow. Figure S6 summarizes the non-uniform refinement (2) and map-quality metrics for the final map reconstruction. Table S3 reports statistics of cryo-EM data collection and validation as well as that of refinement of a full atomic model. The final map resolution was reported as 5.3 Å as multiple rounds of Gold-standard Fourier shell correlation (GS-FSC) validation gave values ranging from 5.27 to 5.35 Å (Figure S6D). In addition, appropriate map sharpening was difficult given the high B-factor (Figure S6B), and the unsharpened map (Figure S5) was used throughout this work for model building and analysis.

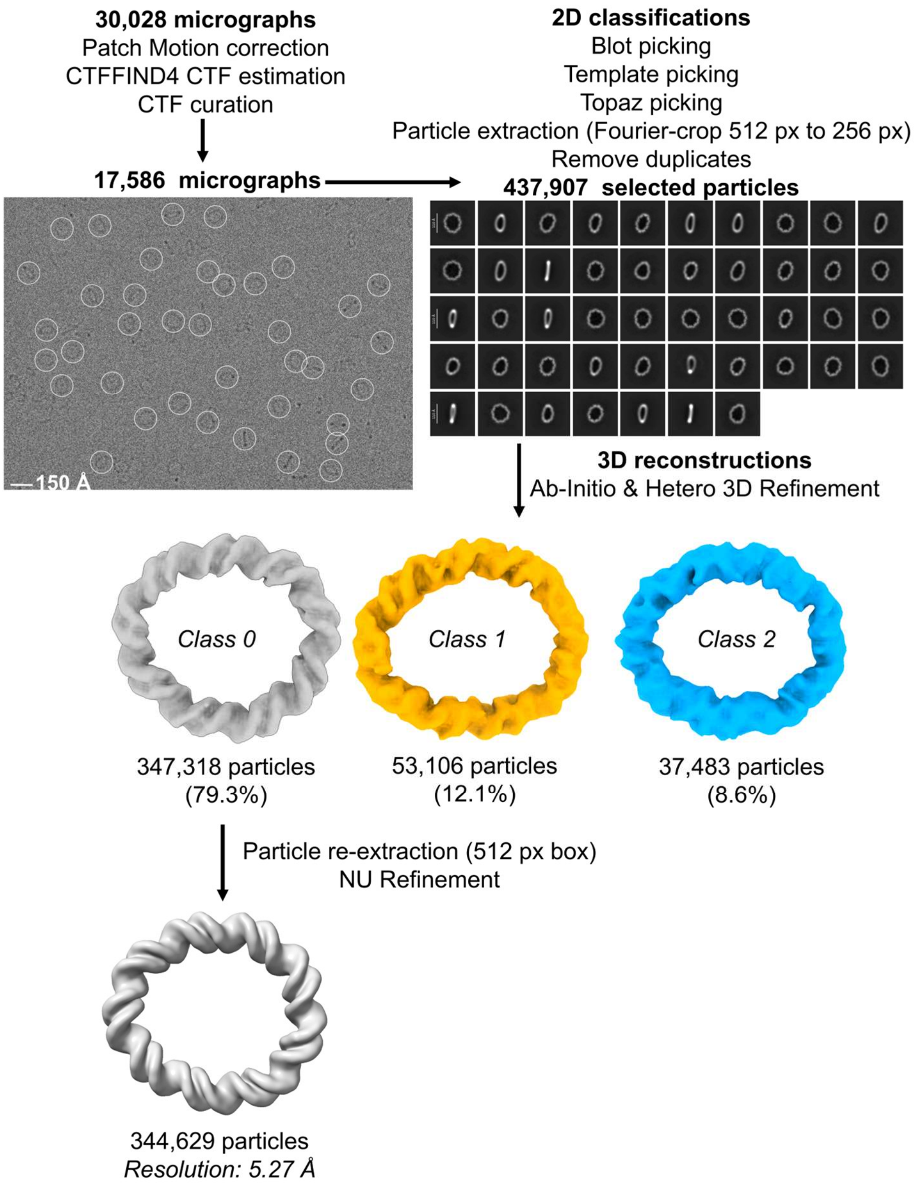

**Figure S5:** Summary of the workflow for dsMC95 cryo-EM data processing.

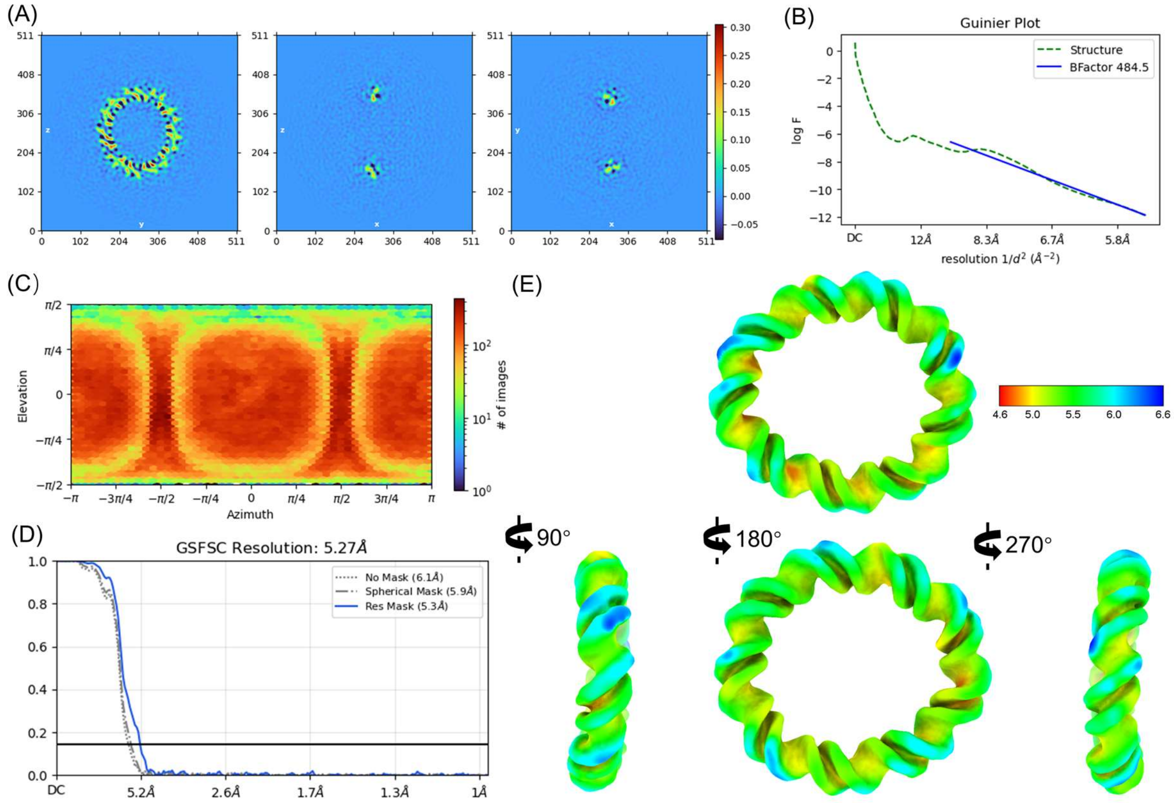

**Figure S6:** Non-uniform refinement results for the Class 0 particles from the dsMC95 dataset. (A) Representative real-space heat-map slices through the refined cryo-EM density. Continuous and well-defined density was observed, indicating that the entire circular DNA molecule was resolved although base-pair level features were absent. (B) Guinier plot showing the overall B-factor estimated from the refined structure. (C) Orientational distribution map of particles used in the reconstruction. (D) Gold-standard Fourier shell correlation (GS-FSC) analysis yielding a global resolution of 5.27 Å. (E) Unsharpened map shown with local resolution estimates. Panels A-D were created in cryoSPARC. Panel E was made in ChimeraX with the local resolution estimated obtained from cryoSPARC.

**Table S3: Cryo-EM data collection, refinement, and validation statistics for dsMC95.**

| | Full atomic model |
|---|---|
| **PDB entry** | ---- |
| **EMDB entry** | ---- |
| **Data collection and processing** | |
| Magnification | 165,000× |
| Voltage (kV) | 300 |
| Electron exposure (e–/Å$^2$) | 50 |
| Defocus range (μm) | -0.5 to -3.0 |
| Pixel size (Å) | 0.51 |
| Symmetry imposed | C1 |
| Initial particle images (no.) | 1,412,668 |
| Final particle images (no.) | 344,629 |
| Map resolution (Å) | 5.3 |
| FSC threshold | 0.143 |
| Map resolution range (Å) | 4.78–6.56 |
| **Refinement** | |
| Initial model used (PDB code) | |
| Model resolution (Å) | 5.4 |
| FSC threshold | 0.143 |
| Model composition | |
| Non-hydrogen atoms | 3,895 |
| Nucleotide | 190 |
| Ligands | 0 |
| *B* factors (Å$^2$) | |
| Nucleotide | 433.61 |
| Ligand | N/A |
| R.m.s. deviations | |
| Bond lengths (Å) | 0.005 |
| Bond angles (°) | 0.790 |
| Validation | |
| MolProbity score | 2.71 |
| Clashscore | 11.18 |
| Poor rotamers (%) | N/A |

#### S3. Additional information on analyzing structural features of dsMC95

##### *S3.1: Analysis of radius*

Radius variations in the full atomic model are shown in Figure 4 of the main text. Figure S7 shows additional information on individual components obtained from simulations.

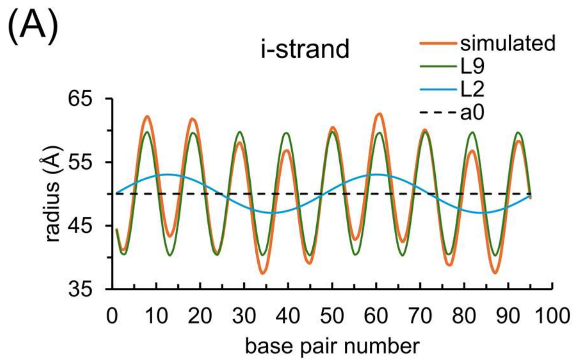

**Figure S7:** Radius analysis of the full atomic model. Overlay of individual components obtained from simulation of radius vs. base-pair-number for the i-strand (A) and the j-strand (B). In each panel, the individual components “a0”, “$L_9$” and “$L_2$” sum to give the “simulated” result. As shown, y-values for L9 and L2 were shifted by the “mid” value to aid comparison. Values of parameters used to generate these fitted curves are reported in the caption of Figure 4B and Figure 4C in the main text.

##### *S3.2. Analysis of groove width*

Figure S8 shows schematics illustrating major and minor groove widths calculations described in Methods in the main text. Table S4 shows examples of major groove width calculations, and Table S5 shows examples of minor groove width calculations.

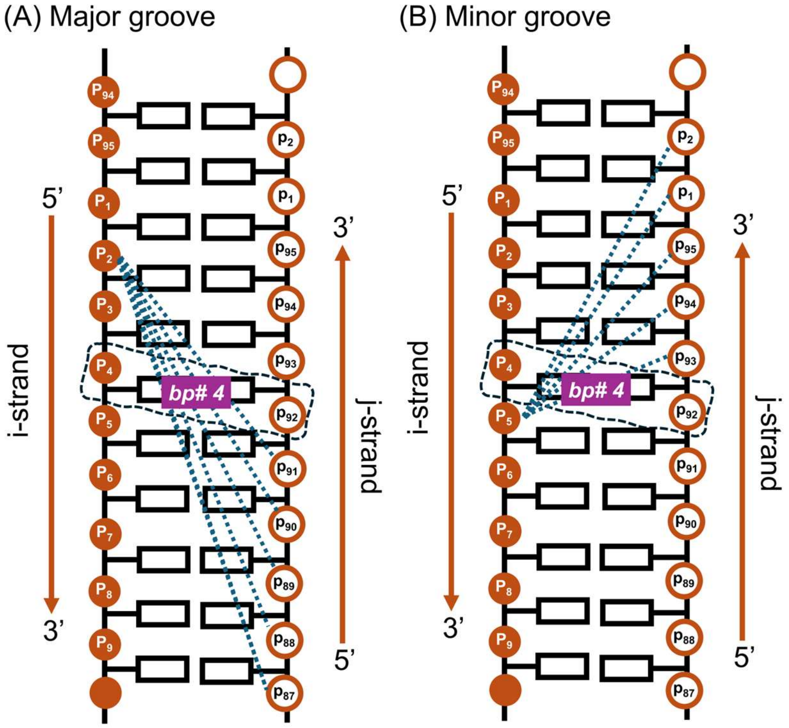

**Figure S8:** Schematics for groove width calculation. In each panel, numbering of the phosphate atoms and the 5’-3’ directionality are marked for the i-strand and the j-strand. As the circular dsMC95 has no termini, at each strand, nucleotide 95 is covalently connected with nucleotide 1 (i.e., $P_{95}$ with $P_1$ in the i-strand and $p_{95}$ with $p_1$ in the j-strand). (A) An example of major groove width calculation at bp#4. The dashed box marks bp#4 that includes $P_4$ of the i-strand and $p_{92}$ of the j-strand. Five distances (dashed lines) were computed between $P_2$ of the i-strand (i.e., $P_{i'}$, i'=m-2=4-2=2, see Methods, main text) and $p_{91}$, $p_{90}$, $p_{89}$, $p_{88}$, and $p_{87}$ of the j-strand (i.e., $p_{j'}$, j'=[95-(m-1)]-n=[95-(4-1)]-n for n=1,2,3,4,5; see Methods, main text). The shortest distance among them was chosen as the major groove width. See also Table S4. (B) An example of minor groove width calculation at bp#4. Five distances (dashed lines) were computed between $P_5$ of the i-strand (i.e., $P_{i''}$, i"=m+1=4+1=5, see Methods, main text) and $p_{93}$, $p_{94}$, $p_{95}$, $p_1$, and $p_2$ of the j-strand (i.e., $p_{j''}$, j"=[95-(m-1)]+n=[95-(4-1)]+n for n=1,2,3; and if j"={[95-(m-1)]+n}-95 for n=4,5; see Methods, main text), and the smallest among them was chosen as the minor groove width. See also Table S5.

Note that a simplified but commonly used approach to calculate major groove width (majGW) is to compute distances between phosphates spanning a fixed number of four base-pairs (3), for example, at bp# m=4, distance between $P_2$ of the i-strand and $p_{91}$ of the j-strand (Figure S8A). However, with the highly bent dsMC95, this approach was found not applicable (see Table S4). Instead, an established procedures (4) were implemented to compute the minimal P-P distance spanning the major groove, which was then set as majGW (Methods, main text). A similar approach was implemented to compute minor groove width (minGW).

**Table S4: Examples of major groove width computation[(a)]**

| bp# = m | i' = m-2 | n | j'=[95-(m-1)]-n | $P_{i'} - p_{j'}$ distance (Å) | majGW (Å) | $R_{major}$ (Å) |
|---|---|---|---|---|---|---|
| 1 | 94[(b)] | 1 | 94 | 18.80[(d)] | 18.80 | 55.28 |
| | | 2 | 93 | 19.77 | -- | |
| | | 3 | 92 | 20.92 | -- | |
| | | 4 | 91 | 24.11 | -- | |
| | | 5 | 90 | 27.30 | -- | |
| 4 | 2 | 1 | 91 | 18.96[(e)] | -- | |
| | | 2 | 90 | 17.21 | -- | |
| | | 3 | 89 | 15.93 | 15.93 | 41.79 |
| | | 4 | 88 | 17.58 | -- | |
| | | 5 | 87 | 22.19 | -- | |
| 62 | 60 | 1 | 33 | 16.93[(d)] | 16.93 | 48.34 |
| | | 2 | 32 | 17.97 | -- | |
| | | 3 | 31 | 20.98 | -- | |
| | | 4 | 30 | 25.55 | -- | |
| | | 5 | 29 | 29.46 | -- | |
| 93 | 91 | 1 | 2 | 18.38[(e)] | -- | |
| | | 2 | 1 | 17.50 | 17.50 | 50.48 |
| | | 3 | 95[(c)] | 20.32 | -- | |
| | | 4 | 94[(c)] | 25.05 | -- | |
| | | 5 | 93[(c)] | 30.07 | -- | |

(a) Computed using full atomic model.

(b) $(m-2) \leq 0$, set $i' = (m-2)+95$.

(c) $[95-(m-1)]-n \leq 0$, set $j' = \{[95-(m-1)]-n\}+95$.

(d) This distance between phosphates spans four base-pairs. Simplified calculation (e.g., "direct P-P distances" from 3DNA (3), http://web.x3dna.org) also gives this majGW value.

(e) This distance between phosphates, which spans four base-pairs (see "bp#4" example in Figure S9A) would have been set as majGW in the simplified calculation (e.g., "direct P-P distances" from 3DNA (3), http://web.x3dna.org). However, it is not the shortest distance spanning the major groove face, and therefore is not accepted as the proper majGW.

**Table S5: Examples of minor groove width computation**[a]

| bp# = m | i" = m+1 | n | j"=[95-(m-1)]+n | $P_{i''} - p_{j''}$ distance (Å) | minGW (Å) | $R_{minor}$ (Å) |
|---|---|---|---|---|---|---|
| 1 | 2 | 1 | 1[b] | 16.15 | -- | |
| | | 2 | 2[b] | 12.78 | -- | |
| | | 3 | 3[b] | 10.44 | 10.44 | 38.34 |
| | | 4 | 4[b] | 11.57 | -- | |
| | | 5 | 5[b] | 16.40 | -- | |
| 4 | 5 | 1 | 93 | 15.58 | -- | |
| | | 2 | 94 | 13.03 | -- | |
| | | 3 | 95 | 12.08 | 12.08 | 51.71 |
| | | 4 | 1[b] | 13.66 | -- | |
| | | 5 | 2[b] | 16.87 | -- | |
| 70 | 71 | 1 | 27 | 15.94 | -- | |
| | | 2 | 28 | 14.11 | 14.11 | 57.70 |
| | | 3 | 29 | 14.51 | -- | |
| | | 4 | 30 | 16.74 | -- | |
| | | 5 | 31 | 21.05 | -- | |
| 95 | 1[c] | 1 | 2 | 16.04 | -- | |
| | | 2 | 3 | 13.20 | -- | |
| | | 3 | 4 | 11.47 | 11.47 | 41.10 |
| | | 4 | 5 | 13.48 | -- | |
| | | 5 | 6 | 18.17 | -- | |

(a) Computed using full atomic model.
(b) [95-(m-1)]+n > 95, set j" = {[95-(m-1)]+n}-95.
(c) (m+1) > 95, set i" = (m+1)-95.

##### *S3.3: Additional information on analysis in the dsMC coordinate*

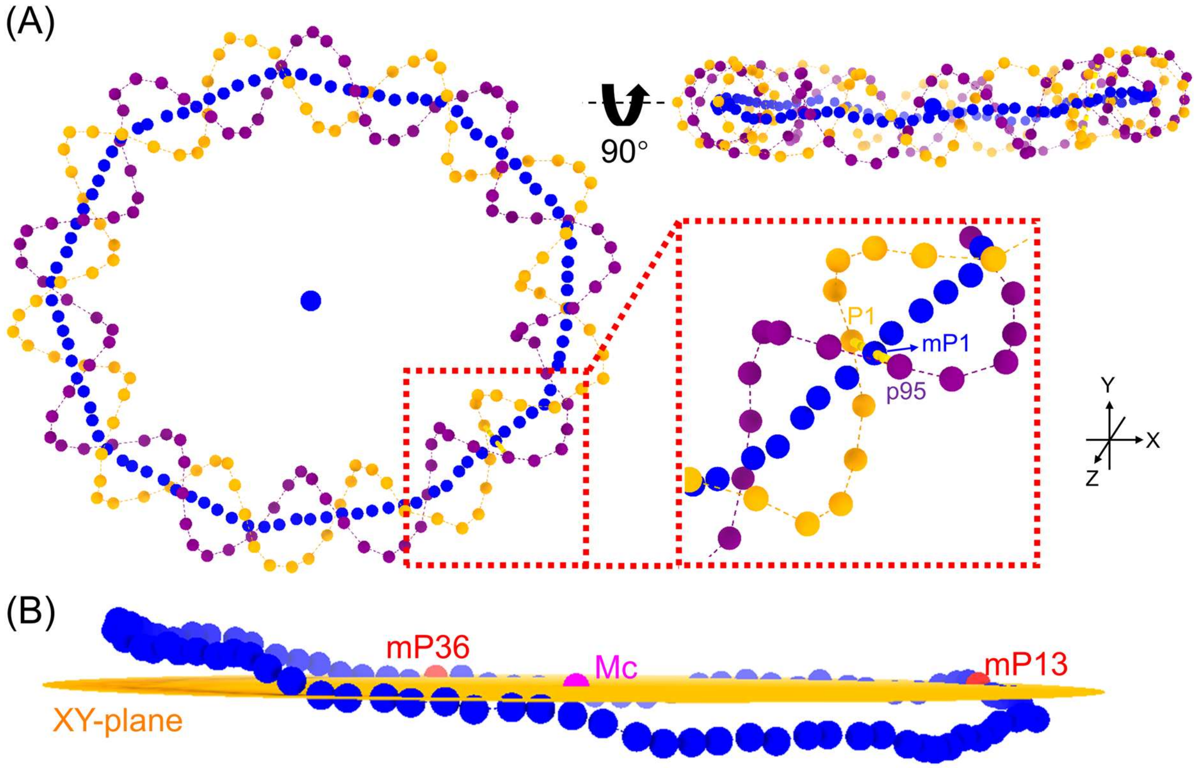

**Figure S9:** mP analysis. (A) mP calculations. Phosphate atoms in the i-strand and j-strand are shown as orange and purple dots, respectively. The computed mP positions are shown as blue dots. The inset shows the mP1 position (130.416, 166.448, 98.551) located at the mid-point of the i-strand $P_1$ atom (138.793, 162.813, 101.504) and the j-strand $p_{95}$ atom (122.038, 170.083, 95.598), which constitute base-pair #1. (B) Out-of-plane displacement analysis. mP positions are shown as blue dots and are viewed perpendicular to the z-axis. The XY-plane of the dsMC coordinate system, defined by Mc (origin), mP13 (+x-axis), and mP36 (+y-axis), is shown in orange.

Examples of calculation of mid-phosphate (mP) points are shown in Figure S9A. Analysis of out-of-plane displacement is shown in Figure S9B.

##### *S3.4. Distortion of the DNA ring observed directly from the cryo-EM map*

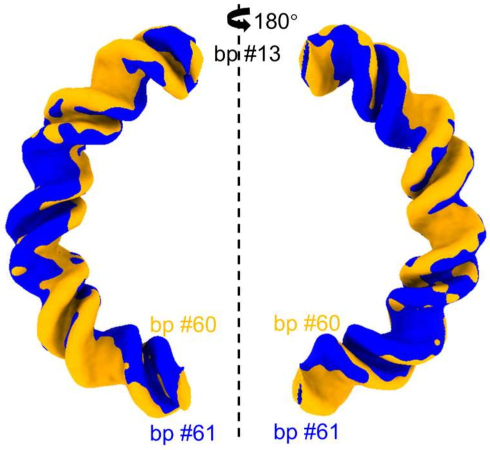

**Figure S10. Analyzing DNA ring distortion.** Unrestricted overlay of cryo-EM maps (contour level 0.018) between the half-circle fragments are shown, with Segment I (bp#13-bp#36-bp#60) in orange and segment II (bp#13-bp#84-bp#61) in blue.

Figure S10 shows un-restricted alignment between the two half-fragments of dsMC95 map, which gives a CC value of 0.97.

#### S4. Additional information on comparison of dsMC95 with other DNA

##### *S4.1. Additional information on comparison with free DNA [T4A4] and [A4T4]*

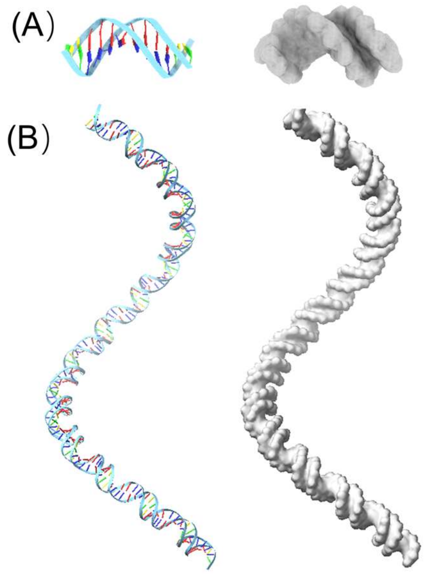

**Figure S11. Free [T4A4] DNA.** (A) The individual [T4A4] DNA (PDB ID 1RVH). Left: atomic model represented in stick and ribbon mode. Right: The corresponding 5Å density map. (B) Model of a d(CT4A4G)$_{15}$ DNA with 15-repeating [T4A4] units. Left: atomic model represented in stick and ribbon mode. Right: The corresponding 5Å density map. The dimensions of the map measured using ChimeraX are 323.90 × 127.94 × 129.36 Å, which are consistent with those reported in reference (7).

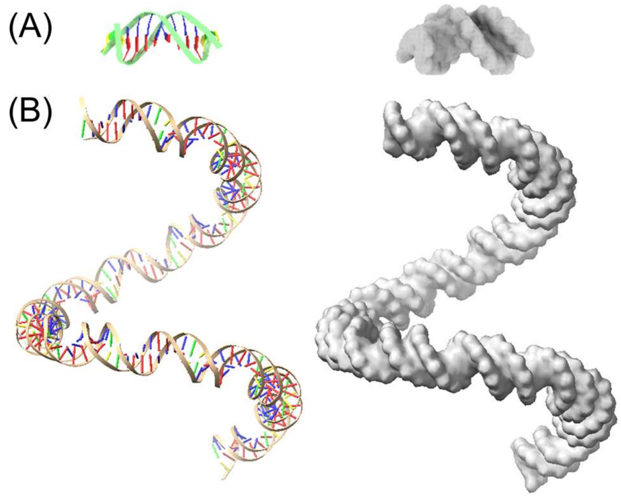

**Figure S12. Free [A4T4] DNA.** (A) The individual [A4T4] DNA (PDB ID 1RVI). Left: atomic model represented in stick and ribbon mode. Right: The corresponding 5Å density map. (B) Model of a d(CA4T4G)$_{15}$ DNA with 15-repeating [A4T4] units. Left: atomic model represented in stick and ribbon mode. Right: The corresponding 5Å density map. The dimensions of the map are 191.41 × 136.98 × 126.54 Å, which are consistent with those reported in reference (7).

Figure S11A shows rendering of the free [T4A4] DNA structure (PDB ID 1RVH (5)) into a 5Å density map in ChimeraX using the molmap program. Following a previously reported procedure (5), local helical parameters from the central 10 base pairs of the [CT4A4G] structure were extracted using 3DNA (3). The parameters were then used to generate an atomic model of a d(CT4A4G)$_{15}$ duplex containing 15-repeating [T4A4] units (Figure S11B, left). The atomic model was then rendered into a 5Å density map (Figure S11B, right). The same approach was applied to generate 5Å density maps of an individual free [A4T4] DNA (PDB ID 1RVI (5)) (Figure S12A) and a d(CA4T4G)$_{15}$ duplex containing 15 repeating [A4T4] units (Figure S12B).

Figure S13 and S14 show overlay between the dsMC95 map with the 5Å maps of individual [T4A4] and [A4T4] DNA, respectively. The maximum CC obtained was 0.85 for [T4A4] (Figure

S13) and 0.91 for [A4T4] (Figure S14), indicating that [A4T4] is more similar to dsMC95.

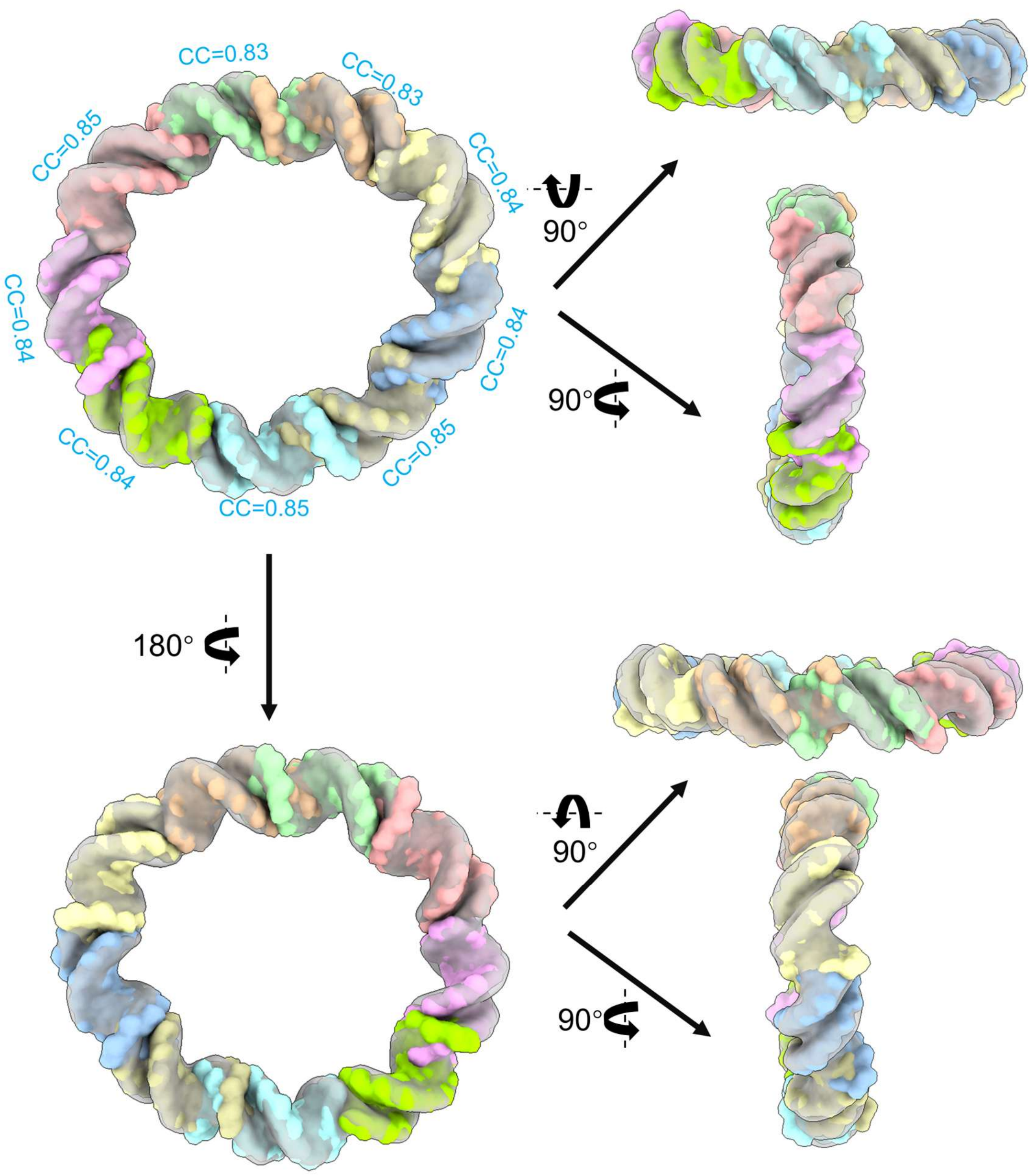

**Figure S13.** Alignment of the individual [T4A4] DNA with dsMC95. dsMC95 is shown in gray, and 5Å synthetic maps of the individual [T4A4] DNA (Figure S11A) are shown in color. CC values are listed next to each corresponding overlay.

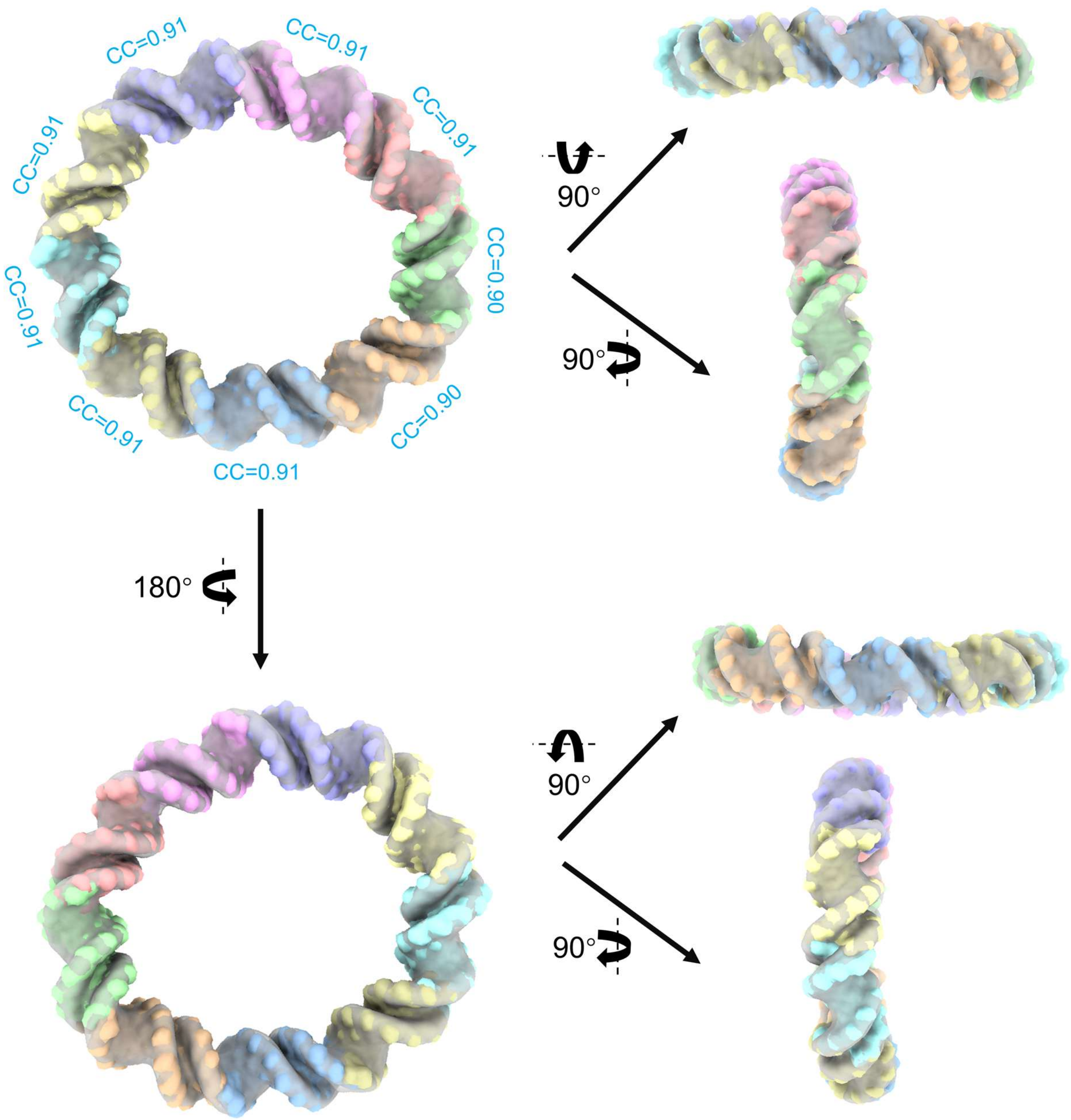

**Figure S14.** Alignment of the individual [A4T4] DNA with dsMC95. dsMC95 is shown in gray, and 5Å synthetic maps of the individual [A4T4] DNA (Figure S12A) are shown in color. CC values are listed next to each corresponding overlay.

##### *S4.2. Comparison of dsMC95 with nucleosomal DNA*

*S4.2.1. Additional information on comparison between dsMC95 and nucleosomal DNA in structure 1KX5*

The crystal structure of the canonical nucleosome core particle, PDB ID 1KX5 (6), has been used as a well-established structure for DNA bending within the nucleosome. Figure S15A shows the 5Å density map of the DNA within the 1KX5 structure generated in ChimeraX using the molmap program. The synthetic 1KX5 map was compared with the dsMC95 map (Figure 7, main text). As shown in Figure S15B, the two maps aligned well between a segment spanning approximately 34 base-pairs.

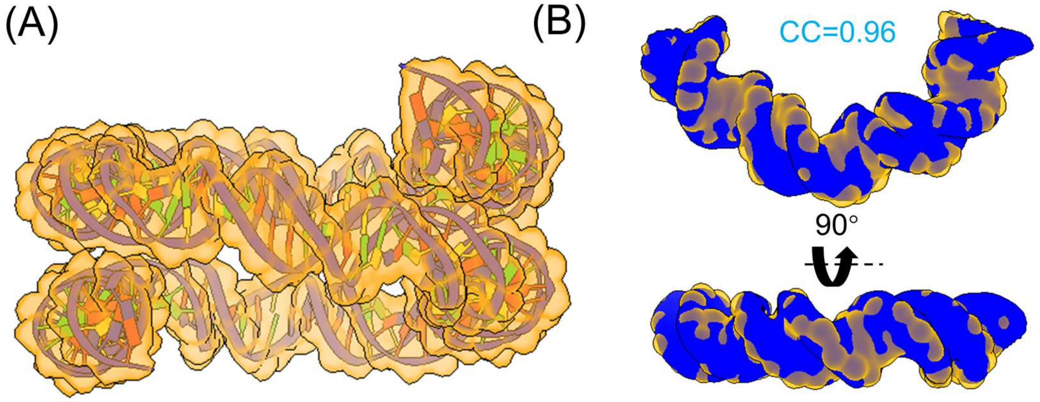

**Figure S15.** Comparison between dsMC95 and nucleosomal DNA from PDB ID 1KX5. (A) DNA structure in PDB ID 1KX5 (stick-ribbon) rendered as a 5Å density map using Chimera-X (orange contour). (B) Overlay between the most similar segments of dsMC95 (blue) and 1KX5 DNA (orange). The dsMC95 segment corresponds to bp#93-bp#31.

*S4.2.2. Comparison between dsMC95 and nucleosomal DNA in structure 7OHC*

Following the same procedures described in sect. S4.2.1, another comparison was made between dsMC95 and nucleosomal DNA from PDB ID 7OHC (7), a nucleosome structure obtained using cryo-EM and currently has the highest resolution. As shown in Figure S16, dsMC95 and DNA from 7OHC align well between a segment spanning approximately 35 base-pairs (Figure S16B). The 7OHC DNA adopts a smaller overall diameter in the XY plane and exhibits a larger span along the Z direction (Figure S16C). These features are consistent with those observed in the comparison between dsMC95 and the 1KX5 DNA (Figure 7 in main text and Figure S15).

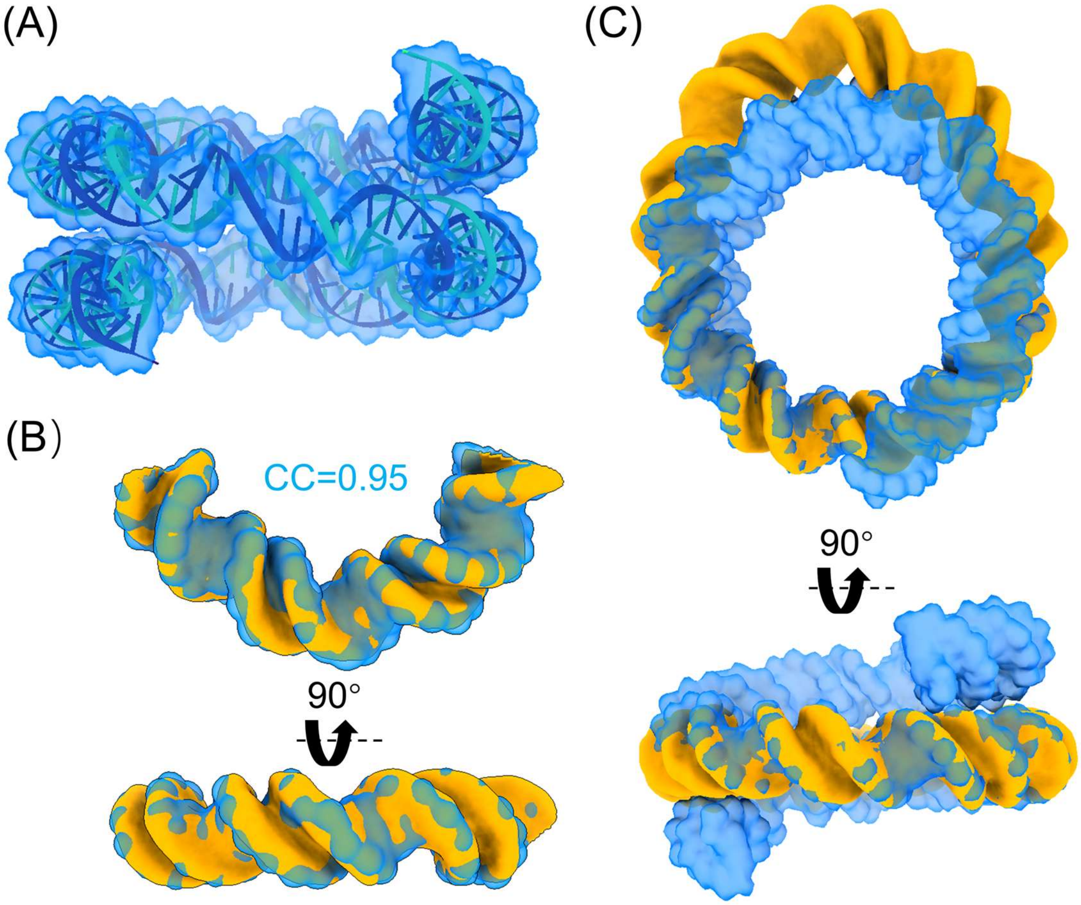

**Figure S16.** Comparison between dsMC95 and nucleosomal DNA from PDB ID 7OHC. (A) DNA structure in PDB ID 7OHC (stick-ribbon) rendered as a 5Å density map using Chimera-X (blue contour). (B) Overlay between the most similar segments of dsMC95 (orange) and 7OHC DNA (blue). The dsMC95 segment corresponds to bp#91-bp#30. (C) Overlay between the entire dsMC95 (orange) and 7OHC DNA (blue).
